# Retroconversion of estrogens into androgens by bacteria *via* a cobalamin-mediated methylation

**DOI:** 10.1101/714501

**Authors:** Po-Hsiang Wang, Yi-Lung Chen, Sean Ting-Shyang Wei, Kan Wu, Tzong-Huei Lee, Tien-Yu Wu, Yin-Ru Chiang

**Author notes:** Author contributions: Y.-R.C. conceptualize this research; P.-H.W., Y.-L.C., K.W. and T.-Y.W. performed experiments; T.-H.L. performed NMR-based structure elucidation; S.T.-S.W performed bioinformatic analysis; P.-H.W., Y.-L.C., and Y.-R.C. analyzed data; and P.-H.W. and Y.-R.C. wrote this manuscript with helps from all the co-authors. Earth-Life Science Institute, Tokyo Institute of Technology, Tokyo, Japan (Po-Hsiang Wang). State Key Laboratory of Marine Environmental Science, Xiamen University, China (Yi-Lung Chen). To whom correspondence should be addressed. Yin-Ru Chiang. Biodiversity Research Center, Academia Sinica, 128 Academia Road Sec. 2, Nankang, Taipei 115, Taiwan.

## Abstract

Steroid estrogens modulate physiology and development of vertebrates. Biosynthesis of C_18_ estrogens from C_19_ androgens by the O_2_-dependent aromatase is thought to be irreversible. Here, we report a denitrifying *Denitratisoma* sp. strain DHT3 capable of catabolizing estrogens or androgens anaerobically. Strain DHT3 genome contains a polycistronic gene cluster *emtABCD* differentially transcribed under estrogen-fed conditions. *emtABCD* encodes a cobalamin-dependent methyltransferase system conserved among estrogen-utilizing anaerobes; *emtA*-disrupted strain DHT3 can catabolize androgens but not estrogens. These data, along with the observed androgen production in estrogen-fed strain DHT3 cultures, indicate the occurrence of a cobalamin-mediated estrogen methylation to form androgens. Consistently, the estrogen conversion into androgens in strain DHT3 cell-extracts requires methylcobalamin and is inhibited by propyl-iodide, a specific inhibitor of cobalamin-dependent enzymes. The identification of the cobalamin-mediated estrogen methylation thus represents an unprecedented metabolic link between cobalamin and steroid metabolism and suggests that retroconversion of estrogens into androgens occurs in the biosphere.

## Introduction

Sex steroids, namely androgens (*e.g.*, testosterone and dihydrotestosterone) and estrogens (*e.g.*, estrone and estradiol), modulate physiology, development, and reproduction of animals (1–4). Biosynthesis of C_18_ estrogens from C_19_ androgens proceeds through the removal of the C-19 angular methyl group, resulting in the formation of an aromatic A-ring (5). This aromatization proceeds through two consecutive hydroxylations of the C-19 methyl group and subsequent oxidative bond cleavage between steroidal C-10 and C-19, which is catalyzed by an aromatase (namely P450arom or CYP19) at the cost of three NADPH and three O_2_ (6). The reverse reaction (from estrogens to androgens) is thermodynamically challenging and has not been reported in any organisms.

Sex steroids are exclusively *de novo* synthesized by eukaryotes; nevertheless, bacteria are major consumers of steroids in the biosphere (7). Interestingly, recent studies suggested that sex steroids mediate bidirectional interactions between bacteria and their eukaryotic hosts (8, 9). Meanwhile, bacteria can also alter a host’s sex steroid profile (10). For example, intestinal *Clostridium scindens* is capable of converting glucocorticoids into androgens (11); *Comamonas testosteroni*, an opportunistic human pathogen, is capable of using a host’s androgens as the sole carbon source and electron donor (12). Furthermore, an earlier study of fecal microbiome suggested that the phylogenetic profile of gut microbiota likely affects endogenous estrogen metabolism in postmenopausal women (13).

Biochemical mechanisms involved in bacterial androgen catabolism have been studied extensively, which includes an O_2_-dependent 9,10-*seco* pathway and an O_2_-independent 2,3-*seco* pathway (14–18). In contrast, current knowledge of the mechanisms involved in estrogen catabolism is very limited. The low aqueous solubility of estrogens (∼1.5 mg/L at room temperature) (19) and the stable aromatic A-ring render estrogen a difficult substrate. Therefore, aerobic bacteria employ O_2_ as a co-substrate of oxygenases to activate and to cleave the aromatic A-ring through the 4,5-*seco* pathway (20–22). Obviously, anaerobic bacteria should develop an O_2_-independent strategy to overcome the chemical recalcitrance of estrogens. To date, only *Denitratisoma oestradiolicum* and *Steroidobacter denitrificans* have been reported to utilize estrogens under anaerobic conditions (23, 24). However, the biochemical mechanism for the anaerobic estrogen catabolism remains completely unknown.

In this study, we enriched an estrogen-degrading denitrifying bacterium *Denitratisoma* sp. strain DHT3 from a municipal wastewater treatment plant, which exhibits high efficiency in estrogen degradation under denitrifying conditions. We first characterized strain DHT3 and annotated its circular genome. Subsequently, we performed comparative transcriptomic analysis to identify the genes potentially involved in the anaerobic estrogen catabolism. The results along with bridging PCR analysis revealed a polycistronic gene cluster *emtABCD* that is differentially expressed in the strain DHT3 transcriptome under estrogen-fed conditions. Bioinformatic analysis predicted that the *emtABCD* gene cluster encodes a putative cobalamin-dependent methyltransferase, which is also present in *D. oestradiolicum* and *S. denitrificans* but not in other steroid-degrading anaerobes incapable of utilizing estrogens. Moreover, the *emtA*-disrupted strain DHT3, although incapable of growing on estrogens, is capable of growing on androgens. Finally, the ^13^C metabolite profile revealed estrogen consumption followed by androgen production in estradiol-fed strain DHT3 cultures. These data indicate the involvement of an unprecedented estrogen conversion into androgens in strain DHT3. Consistently, estradiol is converted into the androgens in strain DHT3 cell-extracts with methylcobalamin or *S*-adenosyl-methionine (SAM) as the methyl donor. Furthermore, the androgen production is significantly inhibited by propyl-iodide, a specific inhibitor of cobalamin-dependent enzymes, in a reversible manner with daylight.

## Results and Discussion

### Enrichment and characterization of *Denitratisoma* sp. strain DHT3

The estrogen-degrading mixed culture was enriched from a denitrifying sludge that was collected from the Dihua Sewage Treatment Plant (Taipei, Taiwan). The estrogen-degrading denitrifier in the enrichment culture was highly enriched by repeating 10^-8^ dilution transfers in a chemical-defined mineral medium containing estradiol as the sole substrate and nitrate as the terminal electron acceptor until a microscopically pure enrichment culture (vibrio-shaped cells) was obtained (**Fig. S1**). Growth of the estradiol-degrading denitrifier on solid media (agar or gelrite plate) was not observed. The phylogenetic analysis showed that the estradiol-degrading denitrifier shares a 97.5 % 16S rRNA gene similarity with *D. oestradiolicum* DSM 16959, suggesting that it belongs to the genus *Denitratisoma*. Therefore, the estradiol-degrading denitrifier is named as *Denitratisoma* sp. strain DHT3 in this study.

Strain DHT3 is able to utilize estradiol, estrone, testosterone, acetate, fumarate, glycerol, and hexanoate as the sole substrate, but was not able to utilize other steroids including synthetic estrogen 17α-ethynyl-estradiol, cholic acid, or cholesterol. The doubling time of strain DHT3 when it grows on estrogens and testosterone ranged from 10–14 hours and 8–10 hours, respectively. No denitrifying growth with the following substrates was observed: yeast extract, peptone, formate, laureate, oleate, pimelate, 2-propanol, butanol, cyclohexanol, cyclopentanone, citrate, glutamate, glucose, fructose, sucrose, benzoate, toluene, or phenol. Moreover, strain DHT3 cannot utilize Fe^3+^, sulfate, O_2_, or perchlorate as the alternative electron acceptor to degrade estradiol.

Stoichiometric analysis suggested that estradiol was mineralized to CO_2_ during the denitrifying growth of strain DHT3 (Fig. 1A). Strain DHT3’s growth (measured based on the increasing protein concentration in culture over time) was in parallel to the consumption of estradiol (electron donor) and nitrate (electron acceptor) in culture (1 L). After 72 hours of incubation, ∼0.5 g estradiol (∼1.8 mmol) and 1.3 g of nitrate (∼21 mM) were consumed, along with ∼380 mg dry cell mass was produced in the 1-L bacterial culture. The complete oxidation of estradiol with nitrate follows the dissimilation equation (24): C_18_H_24_O_2_ + 23NO_3_^-^ + 23H^+^→18CO_2_ + 11.5N_2_ + 23.5H_2_O. Hence, based on nitrate consumption, approximately 0.25 g of estradiol (∼0.9 mmol) was completely oxidized to CO_2_, leaving ∼0.25 g estradiol (∼0.9 mmol) of estradiol to be assimilated into biomass. The amount of assimilated carbon from 0.9 mmol estradiol amounts for ∼195 mg carbon. Assuming that carbon constitutes 50% of dry cell mass (25), the calculated dry cell mass produced from estradiol should be ∼390 mg. This value is close to the observed cell yield (380 mg).

**Fig. 1.**
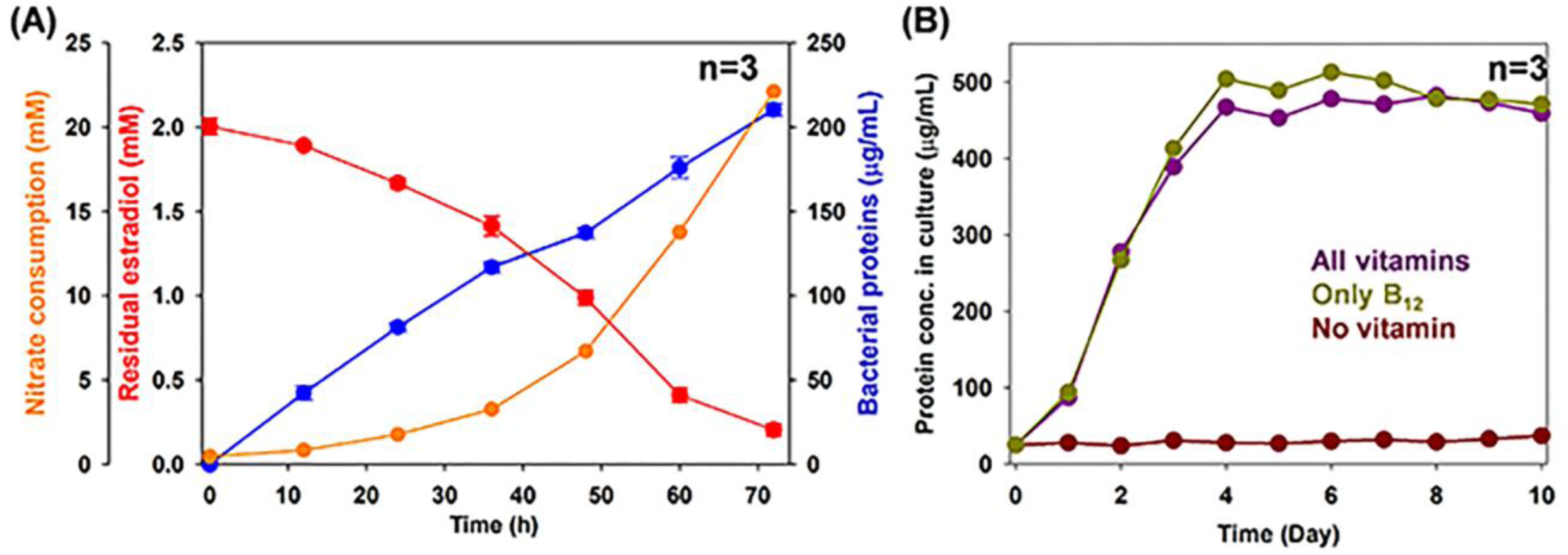
Anaerobic growth of *Denitratisoma* sp. strain DHT3 with estradiol under denitrifying conditions and under different vitamin-supplementing conditions. (**A**) Anaerobic growth of strain DHT3 using estradiol as the sole substrate under denitrifying conditions. (**B**) Anaerobic growth of strain DHT3 on estradiol in the medium supplemented with different vitamins or without vitamins. Bacterial growth was measured based on the increasing total protein concentrations in the cultures. Data shown are means ± SE of three experimental measurements.

We also tested the vitamin requirements of strain DHT3 during denitrifying growth under following conditions: without any vitamins, with an individual vitamin (cyanocobalamin, biotin, calcium pantothenate, thiamine, *p*-aminobenzoic acid, nicotinic acid, or pyridoxamine), and with the VL-7 vitamin mixture composed of the 7 vitamins (26). The growth experiments showed that cobalamin (namely vitamin B_12_; 20 μg/L) is required for strain DHT3’s growth (Fig. 1B). Addition of other vitamins did not facilitate strain DHT3 growth on estradiol (**Fig. S2**), suggesting that strain DHT3 is cobalamin-auxotrophic.

### Identification of estrogen catabolic genes in strain DHT3

We first analyzed the strain DHT3 genome to identify genes potentially involved in the anaerobic estrogen catabolism (**Dataset S1**). The circular genome of strain DHT3 (3.66 Mb; 64.9% G + C; accession CP020914) has been sequenced and annotated (27). Consistent with the observed phenotype, the strain DHT3 genome contains a complete set of androgen catabolic genes in the established 2,3-*seco* pathway, including the genes involved in steroidal A/B-ring degradation (B9N43_01910 to 1920) and C/D-ring degradation (B9N43_4420 to 4465) (Fig. 2A). We also found the 17β-hydroxysteroid dehydrogenase gene (B9N43_02940) responsible for the NAD^+^-dependent dehydrogenation of estradiol to form estrone. Moreover, the strain DHT3 genome lacks most genes for anaerobic cobalamin biosynthesis (28, 29), while it possesses the genes for cobalamin transport (B9N43_10265 to 10280) and utilization (B9N43_10425 to 10440) (Fig. 2A).

**Fig. 2.**
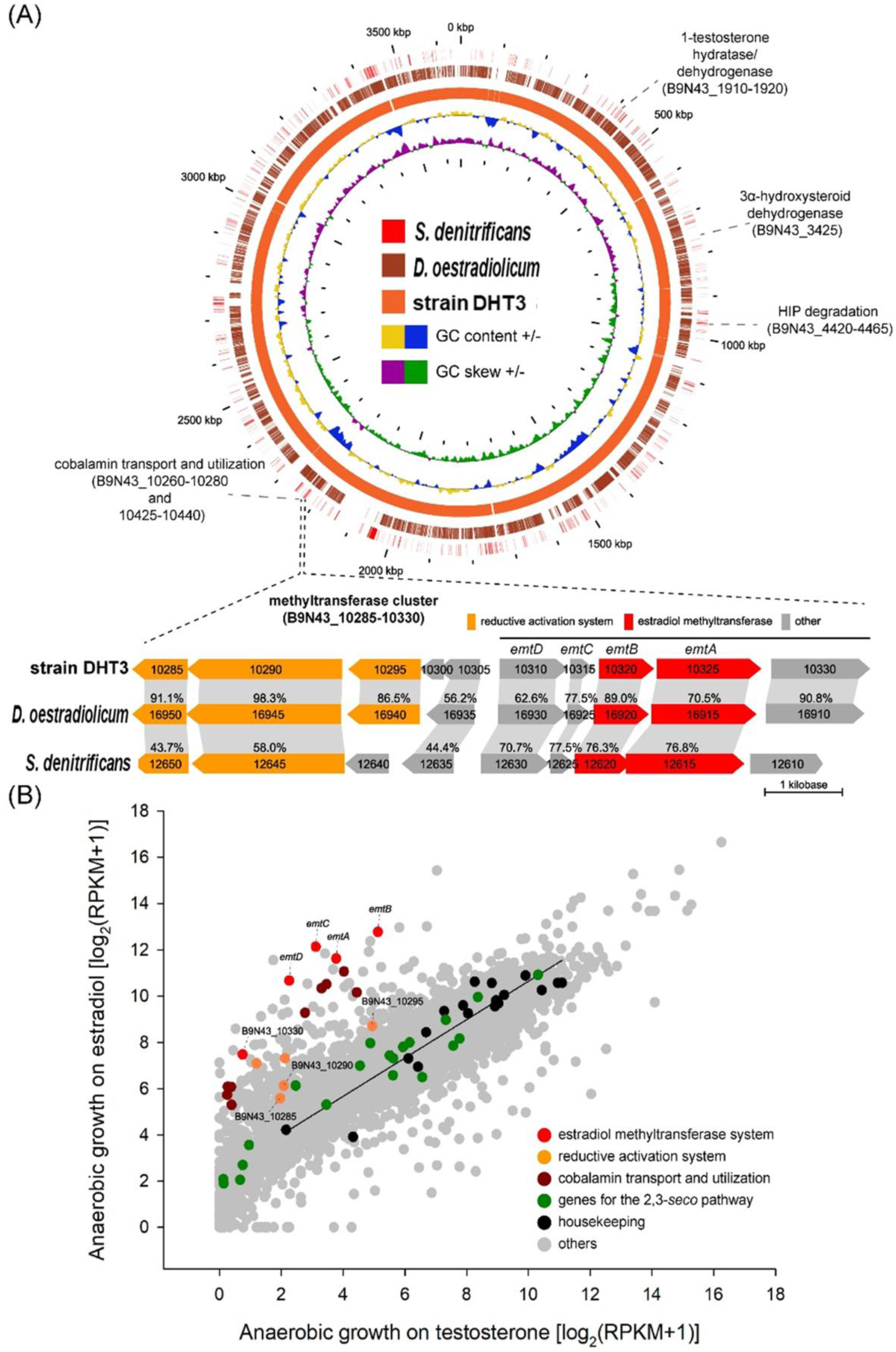
C**o**mparative **genomic analysis and comparative transcriptomic analysis of *Denitratisoma* sp. strain DHT3.** (**A**) Steroid catabolic genes and the putative estrogen catabolic genes on circular genomes of strain DHT3, *D. oestradiolicum* DSM 16959, and *S. denitrificans* DSM 18526. The gene cluster *emtABCD* encoding putative estradiol methyltransferase is polycistronically transcribed in strain DHT3 and is present in these three estrogen-degrading anaerobes. Homologous open reading frames (colored arrows) between different bacterial genomes are connected with gray-colored blocks. Percentage (%) indicate the shared identity of the deduced amino acid sequences. (**B**) Global gene expression profiles (RNA-Seq) of strain DHT3 anaerobically grown on estradiol or testosterone. Each spot represents a gene. The linear regression line is based on the data points of the selected housekeeping genes.

Subsequently, we performed a comparative transcriptomic analysis to detect the genes up-regulated in the estradiol-fed cultures but not in the testosterone-fed cultures. Our data suggested that genes involved in the 2,3-*seco* pathway are commonly expressed (< 4-fold difference) in both estradiol-fed and testosterone-fed cultures (Fig. 2B). In contrast, the genes involved in transport (B9N43_10425 to 10440), salvage (B9N43_10265 to 10280) and reductive activation (B9N43_10285 to 10305) of cobalamin are differentially up-regulated (> 5-fold difference) in the estradiol-fed cultures (Fig. 2B; **Dataset S1**). Among them, B9N43_10285 and _10290 encode a putative methyltransferase-activating protein (30) and a RamA-like ferredoxin (31), respectively. Additionally, a gene cluster putatively encoding a methyltransferase system (B9N43_10310 to 10325; denoted as *emtABCD*) was only up-regulated (> 5-fold difference) in the estradiol-fed culture. Notably, the *emtABCD* gene cluster is also present in estrogen-degrading anaerobes *D. oestradiolicum* and *S. denitrificans* but not in other steroid-degrading bacteria that cannot utilize estrogens (Fig. 2A).

Based on sequence homology (**Dataset S1**), the most EmtA-similar protein in other organisms is MtmB (identity of protein sequence ∼30%), a catalytic subunit of the monomethylamine methyltransferase in methanogenic archaea (32); the most similar protein to EmtB in other organisms is MtmC (identity of protein sequence ∼40%), the corrinoid-binding subunit of the monomethylamine methyltransferase; EmtC is a hypothetical protein; the most similar protein to EmtD is F420/FMN-dependent oxidoreductase (flavodoxin) involved in reductive activation of cobalamin-dependent methyltransferases (33).

We then characterized whether the mRNA products of the *emtABCD* cluster is polycistronically transcribed. Bridging PCR reactions were performed using primers spanning the intergenic regions of these genes (see **Table S1** for individual sequences). The results suggested that B9N43_10310 to 10330 are transcribed polycistronically, including *emtABCD* and B9N43_10330 that encodes a putative serine hydroxymethyltransferase (**Fig. S3**). However, the B9N43_10330-coding protein is less likely a necessary component of the putative cobalamin-dependent methyltransferase since (**a**) the B9N43_10330 homolog is not co-operonic with the *emtABCD* gene cluster in the genome of estrogen-utilizing *S. denitrificans* (Fig. 2A) and (**b**) the expression of B9N43_10330 in both transcriptomes of estradiol-fed and testosterone-fed cultures is significantly lower than that of *emtABCD* (Fig. 2B; **Dataset S1**).

### Functional validation and phylogenetic analysis of *emtA*

Our functional genomic analysis suggested that the polycistronic *emtABCD* gene cluster is likely involved in the anaerobic estrogen catabolism in strain DHT3. Thus, we disrupted the *emtA* gene in strain DHT3 using the TargeTron Gene Knockout System (with a group II intron and the kanamycin-resistant gene inserted) to validate the function of EmtABCD in the anaerobic estrogen catabolism in strain DHT3. We selected *emtA* for the gene disruption experiment since it is annotated as the catalytic subunit of EmtABCD. The *emtA*-disrupted mutant was isolated *via* two successive 10^-8^ dilution transfers in a defined mineral medium with testosterone as the sole substrate and kanamycin (20 μg/mL). PCR with primers flanking the *emtA* gene confirmed successful intragenic insertion of the group II intron into *emtA* in the mutant strain (Fig. 3A). The *emtA*-disrupted strain DHT3 can only utilize testosterone but not estradiol (Figs. 3B **and S4**), revealing that *emtABCD* is involved in the anaerobic estrogen catabolism in strain DHT3.

**Fig. 3.**
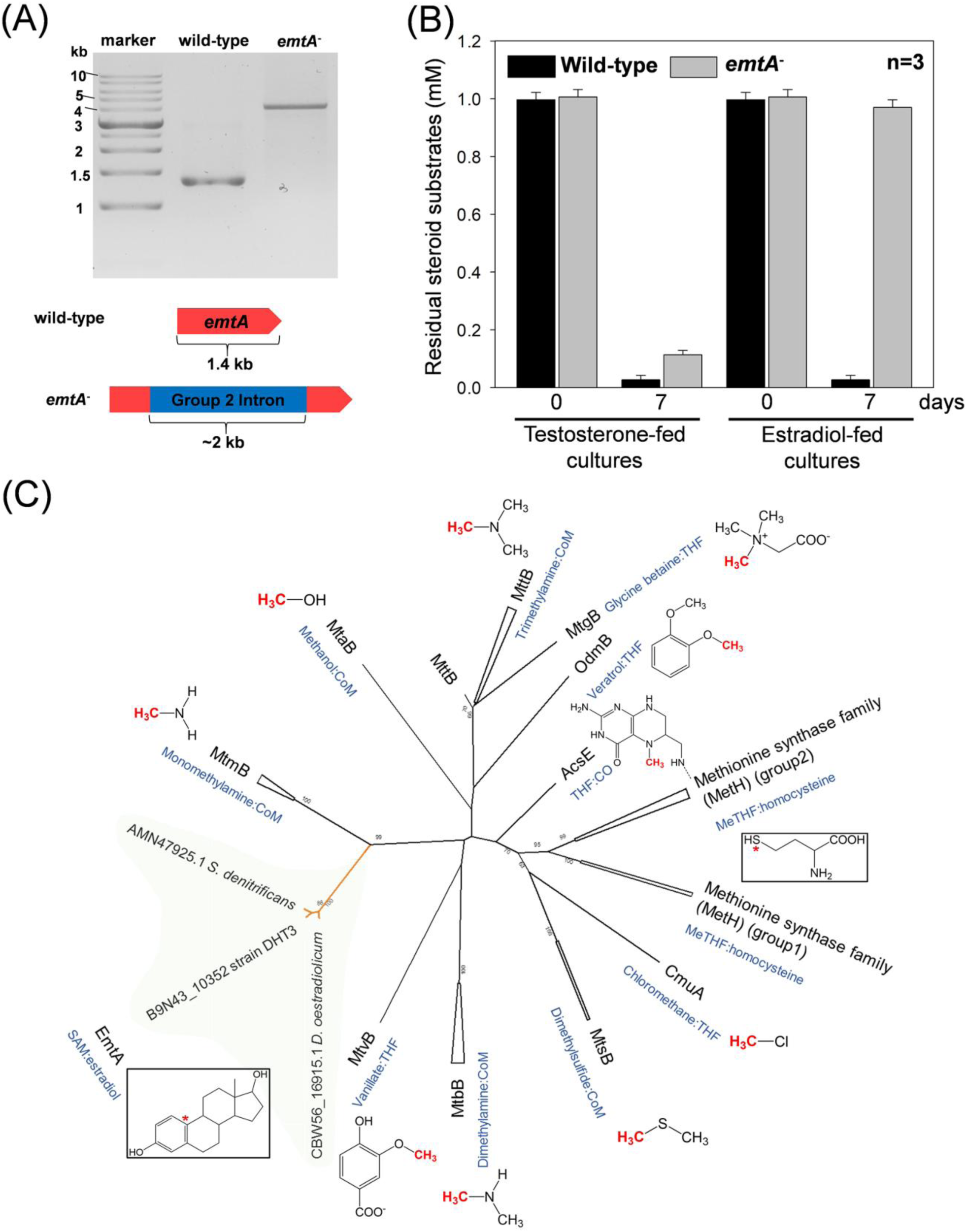
EmtA is involved in the anaerobic estrogen catabolism in *Denitratisoma* sp. strain DHT3. (**A**) Confirmation of intragenic insertion of the group II intron (approximately 2 kb) into the *emtA* of the *emtA*-disrupted mutant (*emtA*^-^) using PCR with the primers flanking this gene. (**B**) Testosterone and estradiol utilizations by the wild-type or *emtA*-disrupted mutant of strain DHT3 (*emtA*^-^). (**C**) Phylogenetic relationship of EmtA and other cobalamin-dependent methyltransferases. The phylogenetic trees were constructed using the neighbor-joining method with Jukes–Cantor parameter and a bootstrap value of 1,000. The star (*) represents the terminal methyl acceptors for MetH and EmtA.

Subsequently, we elucidated the phylogenetic relationship of EmtA and other cobalamin-dependent methyltransferases (**Table S3**). The phylogenetic tree showed that EmtA orthologs from the three estrogen-degrading anaerobes (strain DHT3, *S. denitrificans*, and *D. oestradiolicum*) form a distinct lineage (Fig. 3C), separated from other experimentally characterized cobalamin-dependent methyltransferases in prokaryotes. Moreover, these EmtA orthologs were closely placed into the same clade with the MtmB subunit of monomethylamine methyltransferase in *Methanosarcina* spp. (sequences are shown in **Table S3**), whereas other bacterial cobalamin-dependent methyltransferases were phylogenetically distant from the EmtA orthologs (< 30% sequence similarity) (Fig. 3C). Additionally, pyrrolysine codons, a hallmark of archaeal methylamine:corrinoid methyltransferases (34), are absent in *emtA*. Together, our data reveal that *emtABCD* encoding a putative cobalamin-dependent methyltransferase that is required for the anaerobic estrogen catabolism in strain DHT3.

### Initial step of the anaerobic estrogen catabolism in strain DHT3 proceeds through an unprecedented estrogen conversion into androgens

Next, we managed to identify the estrogen-derived metabolites by analyzing the ^13^C-labelled metabolite profile of the estrogen-fed strain DHT3 cultures (Fig. 4A). The strain DHT3 cultures were anaerobically incubated with a mixture of [3,4C-^13^C]estrone and unlabeled estrone in a 1:1 molar ratio (^13^C-labeled estradiol is not commercially available). The ultra-performance liquid chromatography–high resolution mass spectrometry (UPLC–HRMS) analysis revealed estrone consumption and sequential appearance of several ^13^C-labelled metabolites in the strain DHT3 cultures (Fig. 4B), including six androgenic metabolites (**Table S4**). After 10 hours of incubation, the amounts of the androgenic metabolites in the strain DHT3 cultures significantly decreased with a temporal spike of 17β-hydroxy-1-oxo-2,3-*seco*-androstan-3-oic acid (2,3-SAOA) (Fig. 4) and 3aα-H-4α(3’-propanoate)-7aβ-methylhexahydro-1,5-indanedione (HIP) (**Fig. S5**), two characteristic ring-cleaved intermediates in the 2,3-*seco* pathway (16, 18), revealing that the anaerobic estrogen catabolism in strain DHT3 proceeds *via* an unprecedented C_18_ estrogen conversion into C_19_ androgens. The androgens were further degraded to HIP *via* the established 2,3-*seco* pathway, bringing the consistent expression of genes in the 2,3-*seco* pathway in the estrogen-fed strain DHT3 cultures into context (Fig. 2B).

**Fig. 4.**
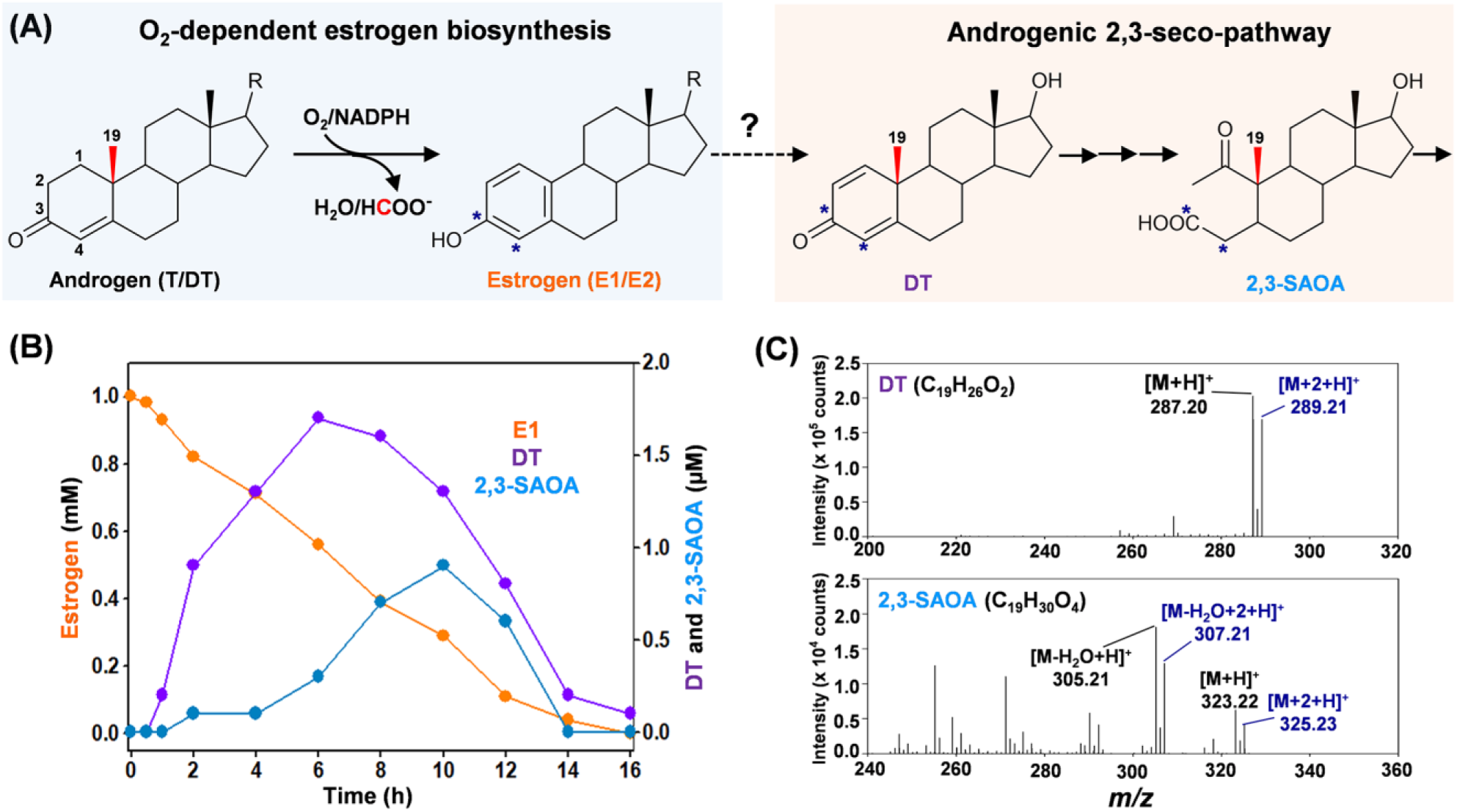
Anaerobic estrogen catabolism by *Denitratisoma* sp. strain DHT3 *via* an unprecedented estrogen conversion into androgens. (**A**) Schematic of estrogen biosynthesis from androgens in eukaryotes and anaerobic estrogen catabolism in strain DHT3 through an unprecedented step of androgen production and subsequent degradation *via* the established 2,3-*seco* pathway. (**B**) Time-dependent estrone (E1) consumption and intermediates production in the strain DHT3 cultures incubated with estrone (1 mM). Data are averages (deviations < 5%) of three experimental measurements. (**C**) UPLC–HRMS-based identification of the androgenic metabolites in the estrone-fed strain DHT3 culture. The estrogen substrate contained unlabeled estrone and [3,4C-^13^C]estrone mixed in a 1:1 molar ratio.

Our data indicate that the observed estrogen conversion into androgens in the strain DHT3 cultures is a methylation reaction likely catalyzed by the putative cobalamin-dependent methyltransferase EmtABCD. Methionine synthase MetH, the best characterized cobalamin-dependent methyltransferase, catalyzes the methyl transfer from 5-methyl-tetrahydrofolate to homocysteine (35). In the primary catalytic cycle (Fig. 5A), the cob(I)alamin prosthetic group of MetH is methylated to form the methylcobalamin using a 5-methyl-tetrahydrofolate as the methyl donor. Subsequently, the methyl group of methylcobalamin is transferred to homocysteine to produce a methionine (36). However, the cob(I)alamin prosthetic group is prone to undergoing single-electron oxidation during the catalytic cycle, yielding the inactive cob(II)alamin even under anaerobic conditions (∼once per 100 turnover) (37); therefore, endergonic reductive activation of the cob(II)alamin prosthetic group is required for the re-entry of MetH to the catalytic cycle (38), which consumes an NADPH (electron donor) and a SAM (methyl donor) for the flavodoxin-mediated methylcobalamin salvage (33,36,39,40) (Fig. 5A). On the other hand, reductive activation of the cob(II)amide prosthetic group in methylamine methyltransferase in methanogenic archaea proceeds through an ATP-dependent mechanism catalyzed by RamA (31) (Fig. 5B). The RamA-like proteins also function in bacteria. For example, a RamA-like ferredoxin Orf7 catalyzes an ATP-dependent reductive activation of cob(II)alamin in *Carboxydothermus hydrogenoformans* (41).

**Fig. 5.**
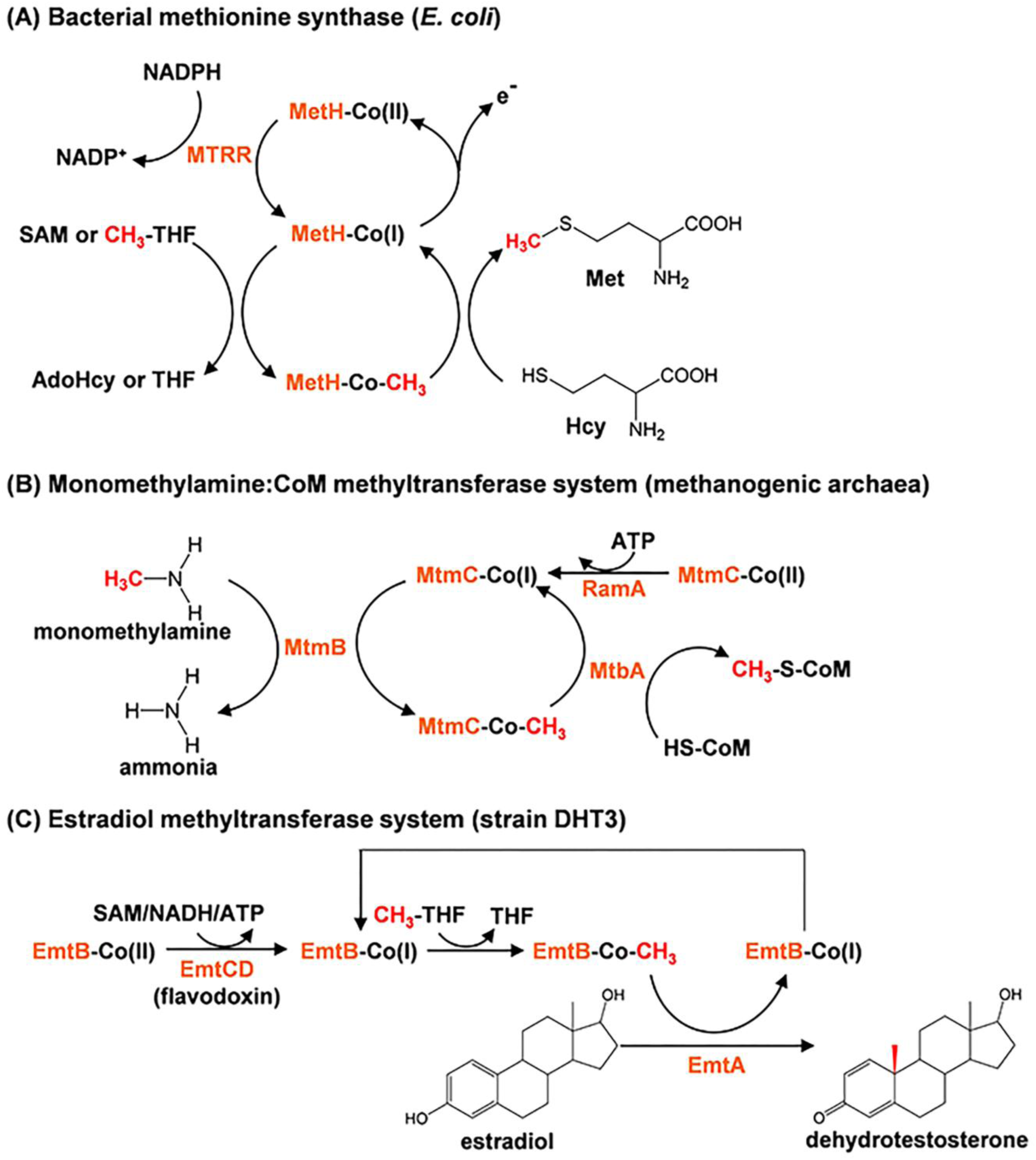
Proposed mechanisms involved in the catalytic cycles of EmtABCD based on the mechanisms of other characterized cobalamin-dependent methyltransferases. (**A)** Methionine synthase MetH: In the catalytic cycle, the cobalamin prosthetic group is methylated by 5-methyl-tetrahydrofolate (CH_3_-THF), followed by the methyl transfer to homocysteine. For reductive activation of the cob(II)alamin prosthetic group, NADH and SAM serve as the electron donor and the methyl donor, respectively. AdoHcy, *S*-adenosylhomocysteine. (**B**) Monomethylamine:CoM methyltransferase MtmBC: the cobamide-binding subunit MtmC forms a heterotrimeric complex with MtbA (the CoM-binding subunit) and MtmB (the catalytic subunit). Reductive activation of the co(II)bamide prosthetic group proceeds through an ATP-dependent reduction catalyzed by RamA. (**C**) Proposed mechanism for the estradiol methylation to form 1-dehydrotestosterone by estradiol methyltransferase EmtABCD in strain DHT3. The cobalamin-binding subunit EmtB and the catalytic subunit EmtA are involved in the catalytic cycle of the estradiol methylation. Reductive activation of the co(II)balamin prosthetic group likely catalyzed by EmtCD (flavodoxin) at the cost of SAM, ATP, or NADH.

### A cobalamin-mediated estradiol methylation to form androgens in strain DHT3

We then managed to validate the occurrence of the cobalamin-mediated estradiol methylation in strain DHT3. Thus, we monitored the methylation of estradiol (mixture of ^2^H-labeled estradiol and unlabeled estradiol in 1:1 molar ratio) in strain DHT3 cell-extracts, along with the addition of the reported methyl donors (SAM and methylcobalamin) and cofactors (cob(I)alamin, NAD(P)H, and ATP) required for the catalysis and reductive activation of cobalamin-dependent methyltransferases (31,39,40,42,43). After incubation, the estradiol-derived metabolites were extracted and analyzed using thin-layer chromatography (TLC) and UPLC–HRMS. We observed the production of two metabolites in the assays containing the strain DHT3 cell-extracts, estradiol, ATP, and methylcobalamin, along with an obvious consumption of estradiol (**Lane 1** in Fig. 6Bi). The UPLC–HRMS analysis indicated that the two estradiol-derived metabolites are both androgens (**Fig. S6**). The ^1^H- and ^13^C-NMR spectra suggest the presence of two hydroxyl groups at C-3 and C-17 as well as two methyl groups (C-18 and C-19) in the steroidal structure, respectively (**Table S5**). The fragment ion profile of the HRMS spectra (**Fig. S6**) and the TLC characteristics of the two androgen metabolites were identical to those of the authentic standards of 17β-hydroxyandrostan-3-one ([M+H]^+^ = 291.23; C_19_H_30_O_2_; denoted as AND1) and 3β,17β-dihydroxyandrostane ([M+H]^+^ = 293.25; C_19_H_32_O_2_; denoted as AND2). The relative abundance of the M1 isotopomers of the two androgenic metabolites in the MS spectra are highly enriched (**Fig. S6**), suggesting that they are downstream metabolites of the ^2^H-labeled estradiol mixture. On the other hand, the androgenic metabolites were not produced in the assays when estradiol, cell-extracts, ATP, or methylcobalamin was excluded (Fig. 6Bi). SAM addition significantly enhanced androgen metabolite production in the strain DHT-3 cell-extracts in a dose-dependent manner (**Fig. S7**). These trends are analogous to the case of reductive activation of the cob(I)alamin prosthetic group in MetH.

**Fig. 6.**
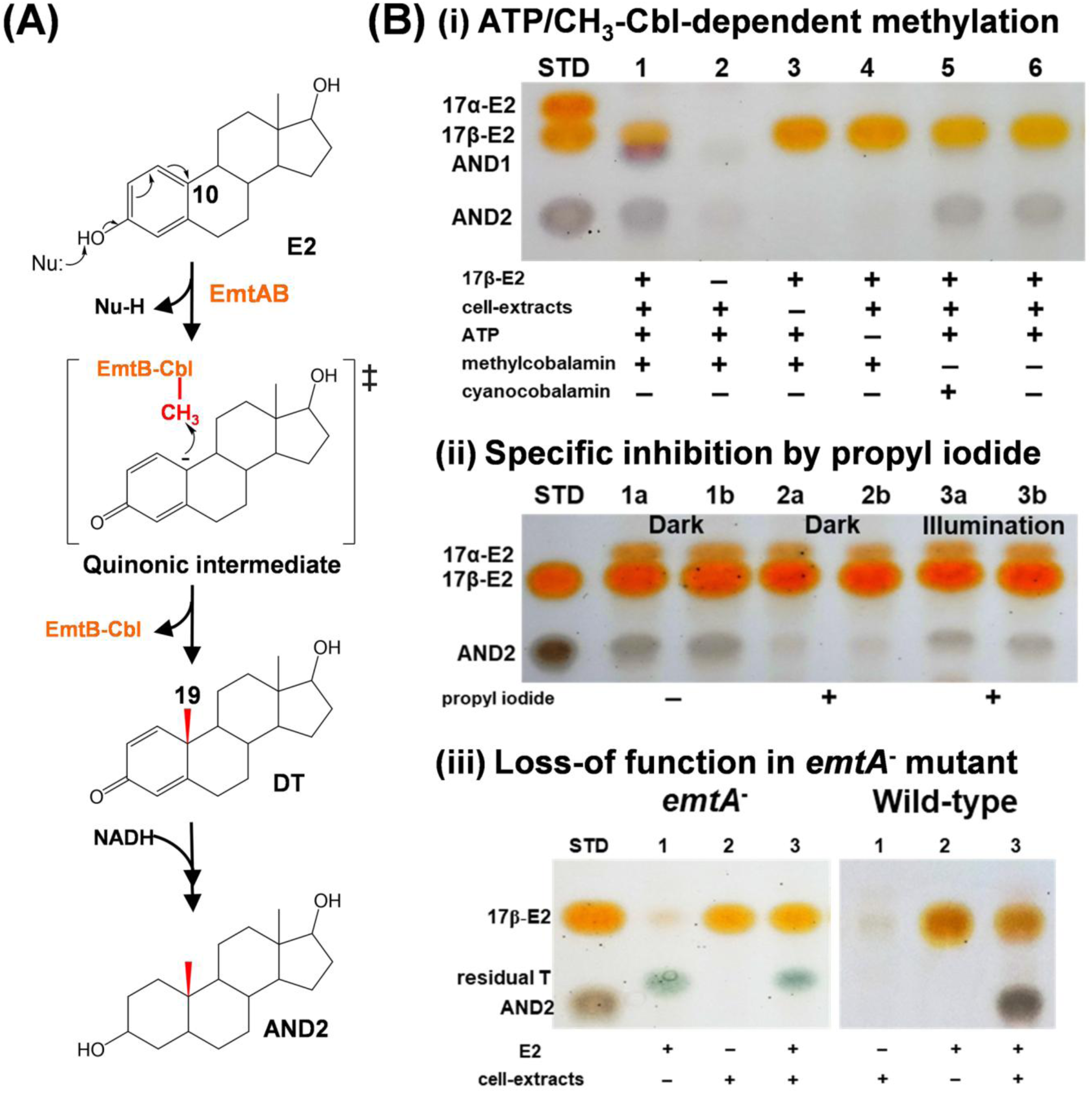
Proposed mechanism for the EmtAB-catalyzed, cobalamin-mediated estradiol methylation. (**A**) Proposed mechanism involved in estrogenic A-ring activation and subsequent cobalamin-mediated C-10 methylation to form androgens. (**B**) Thin-layer chromatographic (TLC) analysis of the cobalamin-mediated estradiol methylation in the strain DHT3 cell-extracts. (**i**) ATP (**Lane 4**) and methylcobalamin (**Lanes 5** and **6**) are required for the estradiol (E2) methylation; (**ii**) specific inhibition of the E2 methylation in the strain DHT3 cell-extracts by propyl-iodide (**Lanes 2a/2b**) in a reversible manner with daylight (**Lanes 3a/3b**). All assays in (**Bii**) contain E2, cell-extracts, ATP, NADH, and with or without propyl-iodide; (**iii**) Loss of E2 methylation activity in the cell-extracts of the *emtA*-disrupted strain DHT3 mutant (**Lane 3**). Assays 3 and 6 in (**Biii**) contain E2, cell-extracts, ATP, NADH, and methylcobalamin. Abbreviations: AND2, 3β,17β-dihydroxyandrostane; T, testosterone.

Next, we managed to validate the involvement of cobalamin-dependent methyltransferase in the estradiol methylation by adding propyl-iodide, a specific inhibitor for cobalamin-dependent enzymes, to the assays. Propyl-iodide inactivates cobalamin-dependent methyltransferases by propylating the cob(I)alamin prosthetic group in the dark (44, 45). Nevertheless, cobalamin-dependent methyltransferases can regain activity upon exposure to daylight (46). Consistently, propyl-iodide addition significantly inhibited the production of the androgenic metabolites in the assays; the inhibition by propyl-iodide was much less effective in the daylight-exposed assays (Fig. 6Bii). Furthermore, the addition of exogenous methylcobalamin, estradiol, NADH, and ATP to the cell extracts of the *emtA*-disrupted strain DHT3 cultures did not result in the androgen production (Fig. 6Biii), as opposed to the androgen production in the assays added with wild-type strain DHT3 cell-extracts. Altogether, our data confirm that the anaerobic estradiol conversion into androgens in strain DHT3 is a cobalamin-mediated methylation reaction catalyzed by EmtABCD.

In this study, we found that strain DHT3 converts estrogens into androgens through a cobalamin-mediated methylation at C-10 on estrogens (Fig. 5C). However, activation of the phenolic A-ring of estrogens is a prerequisite for this methylation reaction. Given that the activated methyl group (CH_3_^+^) of methylcobalamin serves as an electrophile, the estradiol methylation in strain DHT3 likely includes the formation of a nucleophilic carboanion at C-10 on the phenolic A-ring (Fig. 6A). This catalytic strategy has been reported in many studies of anaerobic aromatic catabolism in denitrifying bacteria (47–51), which first proceeds through the deprotonation of the phenolic hydroxyl group to form a phenolate anion. Subsequently, the lone pair electrons on the deprotonated hydroxyl group are migrated to the *para* carbon atom through resonance of the *pi* system, forming a transient quinonic ring. Similarly, in the case of the estradiol methylation, the formation of a quinonic A-ring would come along with the formation of a C-10 nucleophilic carboanion, enabling the electrophilic attack by the methyl cation (CH_3_^+^) on methylcobalamin, yielding the androgenic products with a quinonic A-ring (Fig. 6A). Consistently, the ^13^C metabolite profile also showed that 1-dehydrotestosterone with a quinonic A-ring was produced in the strain DHT3 cultures following estradiol consumption (Fig. 4B). In the strain DHT3 cell-extracts, the produced 1-dehydrotestosterone was further converted into AND1 and AND2 *via* the reduction of both the C-1 double bond and the C-3 keto group (Fig. 6A) by 3-ketosteroid Δ^1^-reductase and 3β-hydroxysteroid dehydrogenase that are consistently expressed in strain DHT3 cells (**Dataset S1**). This claim is supported by the observed AND2 production in the strain DHT3 cell-extracts with 1-dehydrotestosterone as the substrate and NADH as the reductant (data not shown).

## Conclusion

In this study, we demonstrated that strain DHT3 converts estrogens into androgens *via* a cobalamin-mediated methylation and subsequently catabolizes the androgenic intermediates to HIP through the established 2,3-*seco* pathway. The discovery completes central pathways for bacterial steroid catabolism (Fig. 7). Briefly, anaerobic bacteria utilize a convergent catabolic pathway (the 2,3-*seco* pathway) to catabolize sterols, androgens, and estrogens, while aerobic bacteria adopt divergent pathways to catabolize (**a**) sterols and androgens (9,10-*seco* pathway) and (**b**) estrogens (4,5-*seco* pathway). Nevertheless, all three steroid catabolic pathways finally converge at HIP (Fig. 7) and HIP catabolic genes are conserved in the genomes of all characterized steroid-utilizing bacteria (**Fig. S8**) (7, 52).

**Fig. 7.**
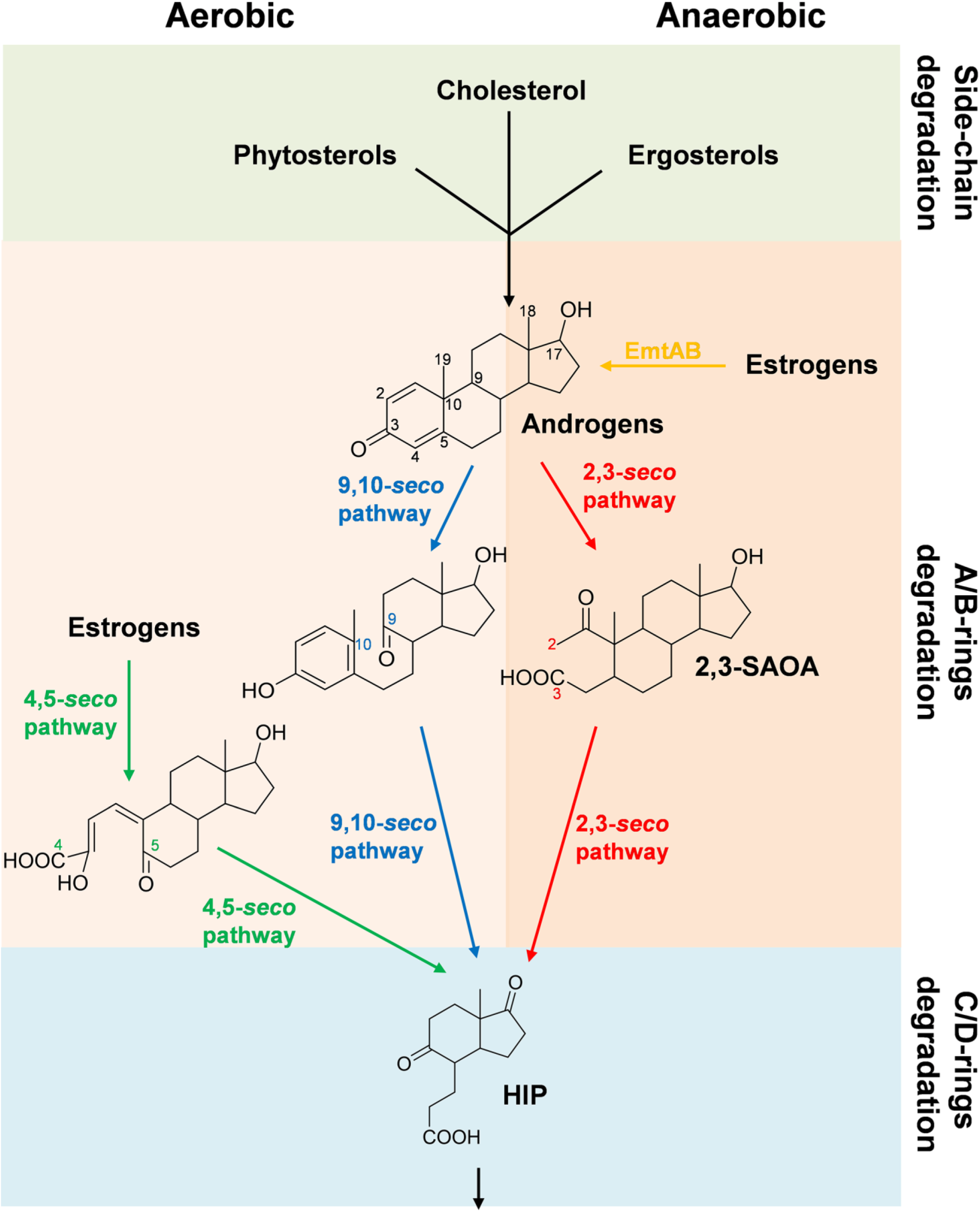
Central pathways for bacterial steroid catabolism. Bacteria adopt a convergent pathway (the 2,3-*seco* pathway) to catabolize different steroids under anaerobic conditions and adopt divergent pathways to catabolize estrogens (the 4,5-*seco* pathway) and other steroids (sterols and androgens; the 9,10-*seco* pathway) under aerobic conditions. All the three steroid catabolic pathways converge at HIP.

Cobamides such as cobalamin are a family of cobalt-containing tetrapyrrole biomolecules with essential biochemical functions in all three domains of life, serving as the prosthetic group for various methyltransferases, isomerases, and reductive dehalogenases (53). Before this study, the known methyl acceptors for cobalamin-dependent methyltransferases included tetrahydrofolate and homocysteine in most organisms (38,42,54) as well as CoM and tetrahydromethanopterin in methanogenic archaea (55). Here, we demonstrated that estrogens are final methyl acceptor of a cobalamin-dependent methyltransferase in a denitrifying proteobacterium, revealing an unprecedented role of cobalamin in steroid metabolism. Furthermore, given that sex steroids are involved in bidirectional metabolic interactions between bacteria and their eukaryotic hosts, finding retroconversion of estrogens into androgens in bacteria portends unexplored microbe-host metabolic interdependencies *via* this newly identified estrogen methylation reaction. Therefore, the *emtABCD* gene cluster can serve as the biomarker to elucidate the occurrence of retroconversion of estrogens in eukaryotic microbiota.

Anaerobic environments such as river and marine sediments are considered as the major reservoirs for estrogens (56). While required by animals, estrogens are also classified as group 1 carcinogens by the World Health Organization (http://monographs.iarc.fr/ENG/Classification/latest_classif.php) and are often detected in surface waters of industrialized countries worldwide (57, 58). Long-term exposure to estrogen-contaminated water at nM levels can disrupt the endocrine system and sexual development in animals (59–61). However, our data reveal that none of the three characterized anaerobic estrogen utilizers possesses complete set of biosynthetic genes to produce cobamide for estrogen utilization. This finding suggests that (**a**) the occurrence of interspecies cobamide transfer between the cobamide producers and the anaerobic estrogen utilizers in natural habitats and (**b**) estrogens are prone to accumulating in O_2_-limited ecosystems devoid of cobamides or cobamide-producing microbes. Therefore, cobalamin or cobalamin-producing anaerobes likely can be augmented into O_2_-limited ecosystems contaminated by estrogens to boost *in situ* bioremediation.

## Materials and Methods

### Chemicals and Bacterial Strains

[3,4C-^13^C]estrone (99%) was purchased from Cambridge Isotope Laboratories (Tewksbury, USA). [16,16,17-D3]17β-estradiol (98 %), 5α-androstan-17β-ol-3-one (=17β-hydroxyandrostan-3-one) and 3aα-H-4α(3’-propanoate)-7aβ-methylhexahydro-1,5-indanedione (HIP) were obtained from Sigma-Aldrich (St. Louis, MO, USA). 5α-androstan-3β,17β-diol (=3β,17β-dihydroxyandrostane) was purchased from Steraloids Inc (Newport, RI, USA). Other chemicals were analytical grade and purchased from Mallinckrodt Baker (Phillipsburg, USA), Merck Millipore (Burlington, USA), and Sigma-Aldrich unless specified otherwise. *Denitratisoma oestradiolicum* strain AcBE2-1 (=DSM 16959) and *Steroidobacter denitrificans* strain FS (=DSM 18526) were purchased from the Deutsche Sammlung für Mikroorganismen und Zellkulturen (DSMZ; Braunschweig, Germany).

### Enrichment of *Denitratisoma* sp. strain DHT3 from an anoxic sludge

Strain DHT3 was enriched from an anoxic sludge collected from the Dihua Sewage Treatment Plant, Taipei, Taiwan (DHSTP; 25°4’20.97”N, 121°30’36.17”E). To enrich the anaerobic estrogen utilizer, the sludge sample (100 mL) was spiked with estradiol (1 mM) and nitrate (10 mM), and was incubated at 28°C under a N_2_/CO_2_ (80:20, v/v) atmosphere in the dark (to avoid growth of phototroph) for 14 days. Later, twenty milliliters of the estradiol-spiked sludge sample was transferred to a chemically defined mineral medium (500 mL) (62) with estradiol (1 mM) as the sole substrate and nitrate as the terminal electron acceptor. The final pH was adjusted to 6.5 with HCl to alleviate the gradual pH increase in culture caused by denitrification. After the added estradiol was mostly consumed, the estradiol-spiked mixed culture (1 mL) was transferred to the same defined medium (200 mL). This step was repeated ten times to selectively enrich the anaerobic estrogen utilizer. The 10^th^ transfer culture was serially diluted from 10^1^ to 10^9^ times in the same defined medium to extinct the non-estrogen degraders, and incubated under the same conditions. After the 10^9^ times-diluted culture consumed most of the added estradiol, the dilution-to-extinction transfers were repeated again. After that, the culture purity was validated microscopically and by growth tests of the enrichment culture in liquid R2A rich medium and in the defined medium added with yeast extracts. The 16S rRNA gene sequence was amplified from the total genomic DNA extracted from the enrichment culture using universal primers 27F and 1492R (63), and the taxonomy of strain DHT3 was determined using BALST. Since the growth of strain DHT3 in the solid medium (agar or gelrite plate) was not observed, we do not exclude the possibility that trace contaminated microbes are present in the microscopically pure strain DHT3 enrichment culture.

### Anaerobic growth of strain DHT3 with and without vitamins

To test the vitamin requirements of strain DHT3, we anaerobically cultivated strain DHT3 in the defined mineral media containing estradiol (3 mM), nitrate (initially 10 mM), and individual vitamins [cyanocobalamin (20 μg/L), biotin (10 μg/L), calcium pantothenate (25 μg/L), thiamine (50 μg/L), *p*-aminobenzoic acid (50 μg/L), nicotinic acid (100 μg/L), or pyridoxamine (250 μg/L)]. The growth of strain DHT3 with and without the mixed vitamin solution VL-7 (26) was served as the positive and negative controls, respectively. The cultures were sampled daily, and the samples (0.5 mL each) were centrifuged at 10,000 × g for 10 min at 4°C. After centrifugation, the bacterial pellets were stored at −20°C before protein content determination. Proteins were extracted from the frozen pellets using B-PER Bacterial Cell Lysis Reagents (Thermo Fisher Scientific; Waltham, MA, USA). The protein content in the samples was estimated using Pierce BCA Protein Assay Kit (Thermo Fisher Scientific) according to manufacturer’s instructions, with a standard curve generated using analytical-grade bovine serum albumin. The nitrate content was measured using Spectroquant Nitrate Test Kit HC707906 (Merck, Germany) according to manufacturer’s instructions.

### Analytical methods

a. **Thin-layer Chromatography (TLC).** The steroid standards and products were separated on silica gel-coated aluminum TLC plates (Silica gel 60 F_254_: thickness, 0.2 mm; 20 × 20 cm; Merck) using dichloromethane:ethyl acetate:ethanol (14:4:0.05, v/v/v) as the developing phase. The steroids were visualized under UV light at 254 nm or by spraying the TLC plates with 30% (v/v) H_2_SO_4_, followed by an incubation for 1 min at 100°C (in an oven).
b. **High-Performance Liquid Chromatography (HPLC).** A reversed-phase Hitachi HPLC system equipped with an analytical RP-C_18_ column (Luna 18(2), 5 μm, 150 × 4.6 mm; Phenomenex) was used for separating steroidal metabolites in this study. The separation was achieved isocratically using a mobile phase of 45% methanol (v/v) at 35°C at a flow rate of 0.5 mL/min. The steroidal metabolites were detected using a photodiode array detector (200–450 nm). In some studies, HPLC was also used for quantifying steroids present in the strain DHT3 cultures. The quantity of steroids was measured using a standard curve generated from individual commercial steroid standards. The *R*^2^ values for the standard curves were at least > 0.98. The presented data are the average values of three experimental measurements.
c. **Ultra-Performance Liquid Chromatography–High Resolution Mass Spectrometry (UPLC–HRMS).** Steroids were analyzed using UPLC**–**HRMS on a UPLC system coupled to either an Electric Spray Ionization**–**Mass Spectrometry (ESI**–**MS) system or an Atmosphere Pressure Chemical Ionization**–**Mass Spectrometry (APCI**–**MS) system. Steroids were firstly separated using a reversed-phase C_18_ column (Acquity UPLC^®^ BEH C18; 1.7 μm; 100 × 2.1 mm; Waters) at a flow rate of 0.4 mL/min at 35°C (oven temperature). The mobile phase comprised a mixture of two solutions: solution A (0.1% formic acid (v/v) in 2% acetonitrile (v/v)) and solution B (0.1% formic acid (v/v) in methanol). Separation was achieved using a gradient of solvent B from 10% to 99% in 8 min. ESI**–**MS analysis was performed using a Thermo Fisher Scientific^TM^ Orbitrap Elite^TM^ Hybrid Ion Trap-Orbitrap Mass Spectrometer (Waltham, MA, USA). Mass spectrometric data in positive ionization mode were collected. The source voltage was set at 3.2 kV; the capillary and source heater temperatures were 360°C and 350°C, respectively; the sheath, auxiliary, and sweep gas flow rates were 30, 15 and 2 arb units, respectively. APCI–MS analysis was performed using a Thermo Fisher Scientific^TM^ Orbitrap Elite^TM^ Hybrid Ion Trap-Orbitrap Mass Spectrometer (Waltham, MA, USA) equipped with a standard APCI source. Mass spectrometric data in positive ionization mode (parent scan range: 50–600 *m/z*) were collected. The capillary and APCI vaporizer temperatures were 120°C and 400°C, respectively; the sheath, auxiliary, and sweep gas flow rates were 40, 5 and 2 arbitrary units, respectively. The source voltage was 6 kV and current 15 μA. The elemental composition of individual adduct ions was predicted using Xcalibur™ Software (Thermo Fisher Scientific).
d. **Nuclear Magnetic Resonance (NMR) spectroscopy.** ^1^H- and ^13^C-NMR spectra were recorded at 27°C using a Bruker AV600_GRC 600MHz NMR. Chemical shifts (δ) were recorded and presented in parts per million with deuterated methanol (99.8%) as the solvent and internal reference.

### General molecular biological methods

The strain DHT3 genomic DNA was extracted using the Presto^TM^ Mini gDNA Bacteria Kit (Geneaid, Taiwan). PCR mixtures (50 μL) contained nuclease-free H_2_O, 2 × PCR buffer (25 μL), dNTPs (2 mM), *Taq* polymerase (1.2 U) (BioTherm; NatuTec GmbH; Frankfurt am Main, Germany), forward and reverse primers (each 200 nM), and template DNA (30 ng). The PCR products were verified using standard TAE-agarose gel (1.5%) electrophoresis with the SYBR® Green I nucleic acid gel stain (Thermo Fisher Scientific), and the PCR products were purified using the GenepHlow Gel/PCR Kit (Geneaid).

### RNA isolation and bridging PCR

Bridging PCR was applied to determine the polycistronism of the *emt* gene cluster. Total RNA was extracted from the estradiol-grown strain DHT3 cells using the Direct-zol RNA MiniPrep Kit (Zymo Research; Tustin, CA, USA). The crude total RNA was further purified using Turbo DNA-free Kit (Thermo Fisher Scientific) to eliminate DNA. The DNA-free total RNA was reversely transcribed to cDNA using the SuperScript^®^ IV First-Strand Synthesis System (Thermo Fisher Scientific). Random hexamer primers were used for cDNA synthesis. The polycistronism of the *emt* gene cluster was analyzed using intergenic PCR with total cDNA as the template and the following PCR program was used: 95°C for 5 min, followed by 25 cycles at 95°C for 30 s, 60°C for 30 s, 72°C for 75 s, and finally 72°C for 3 min. The primers are shown in **Table S1**. The strain DHT3 genomic DNA and total RNA were used as the positive and negative control, respectively.

### RNA extraction and RNA-Seq

**The** strain DHT3 transcriptome was extracted from the strain DHT3 cells anaerobically cultivated with estradiol (2 mM) or testosterone (2 mM) as the sole substrate. The transcriptomes were extracted using the Direct-zol RNA MiniPrep Kit (Zymo Research), and were further purified using Turbo DNA-free Kit (Thermo Fisher Scientific) to eliminate DNA. rRNA was removed from the transcriptome samples using the Ribo-Zero rRNA Removal Kit (Epicentre Biotechnologies; Madison, WI, USA). The quality of the resulting RNA library was assessed using the Experion RNA Analysis Kit (Bio-Rad; Hercules, CA, USA), and only the samples with an integrity number 8-10 were selected for transcriptomic analysis. First-strand cDNA was synthesized using the purified transcriptomes as the templates. Second-strand cDNA was synthesized in a reaction mixture containing Second-Strand Synthesis Kit (New England Biolabs; Ipswich, MA, USA). cDNA was purified using the Qiagen purification kit, followed by end repair using NEBNext^®^ End Repair Module (New England Biolabs). Antisense strand DNA was digested in the samples using the Uracil-Specific Excision Reagent Enzyme Kit (New England Biolabs), followed by a PCR to amplify the cDNA. The constructed sequencing libraries were sequenced as paired-end reads (with 126-bp read length) on the Illumina HiSeq 2000 system (Illumina; San Diego, CA). Adapter of the RNA-Seq reads were removed with cutadapt (Version 1.4.2) and the remaining sequences were trimmed by using Seqtk (Version 1.2-r94). The maximum ambiguous nucleotide number was set to 2; the lasting sequences (min length = 35-bp; error probability < 0.05) were included for subsequent analysis. The constructed sequencing libraries were mapped to the coding DNA sequences in the strain DHT3 genome using Bowtie2 (Version 2.2.3). The mapping results were quantified using eXpress (Version 1.5.1) with software’s default setting and expressed in terms of Reads Per Kilobase of transcript per Million mapped reads (RPKM).

### *emtA* gene disruption in strain DHT3

The *emtA* gene of strain DHT3 was disrupted by an intragenic insertion of a group II intron into this gene using the TargeTron Gene Knockout System Kit (Sigma-Aldrich, St. Louis, MO). The intron insertion sites for *emtA* were predicted using online intron design software (TargeTron gene knockout system; Sigma-Aldrich), and the primer sequences were generated for intron retargeting. The intron PCR template from the TargeTron Gene Knockout System Kit and the following four primers were used for the construction: *emtA*-773|774-IBS, *emtA*-773|774s-EBS1d, *emtA*-773|774s-EBS2, and EBS universal primer (see **Table S1** for individual sequences). The PCR was performed according to the manufacturer’s instructions. The 350-bp PCR product was purified and subsequently ligated into the pACD4K-C Linear Vector. This resulted in a plasmid, pACD4K-C::*emtA*, specific for *emtA* knockout by insertion of the intron fragment between the 773th and 774th base pairs in the sense direction. The construct mixture was firstly transformed into the ECOS^TM^ 101 *E. coli* strain DH5α competent cells via heat-shock incubation (Yeastern Biotech, Taipei, Taiwan). The pACD4K-C::*emtA* purified from the correct clones (sequences verified using Sanger Sequencing) was then transformed to strain DHT3 through electroporation (2,500 V, 25 μF, and 200 Ω). Strain DHT3 does not carry a copy of T7 RNA Polymerase on the chromosome. Therefore, a source of T7 RNA Polymerase was provided by co-transforming another plasmid pAR1219 (Sigma-Aldrich, St. Louis, MO). The plasmids pACD4K-C::*emtA* and pAR1219 were maintained by selection on chloramphenicol and ampicillin, respectively. Strain DHT3 containing the two plasmids was anaerobically incubated in a chemically defined medium containing estradiol (2 mM), chloramphenicol (100 μg/mL) and ampicillin (100 μg/mL). The bacterial culture was cultivated at 28°C under a N_2_/CO_2_ atmosphere (80:20, v/v). When the cultures reached a cell density of ∼8 × 10^8^/mL, isopropyl β-D-1-thiogalactopyranoside (IPTG) (2 mM) was added to the bacterial culture. After overnight incubation, the strain DHT3 cells were harvested through a centrifugation at 3,500 × g for 10 min at 20°C. The *emtA*-disruption was then selected using a kanamycin marker that was activated upon chromosomal insertion. Therefore, the resulting strain DHT3 cell pellet was resuspended and was cultivated in a chemically defined medium containing testosterone (2 mM) and kanamycin (20 μg/mL). Kanamycin-resistant strain DHT3 cells were then isolated by two subsequent serial dilution series (10^1^ to 10^9^ times) using the same medium. Successful *emtA* disruption was confirmed using PCR with primers flanking the *emtA* gene (see **Table S1** for individual sequences).

### Analysis of the steroidal metabolite profile of the estrogen-grown strain DHT3

Strain DHT3 cells in the exponential growth phase (with a cell density of approximately 8 × 10^8^/mL) were harvested through a centrifugation at 8,000 × g for 10 min at 25 °C. After decanting the supernatants, the cell pellets were immediately re-suspended in the same chemically defined medium. The concentrated strain DHT3 suspensions (5 × 10^9^/mL; 10 mL) were spiked with 1 mM of estrone (unlabeled estrone and [3,4C-^13^C]estrone mixed in a 1:1 molar ratio) and was incubated under a N_2_/CO_2_ (80:20, v/v) atmosphere at 28°C with continuous stirring. 17α-ethinylestradiol (final concentration 50 μM), which cannot be utilized by strain DHT3, was added to the concentrated strain DHT3 cell suspensions to serve as an internal control. The cell suspension samples (0.5 mL) were withdrawn every 30–120 min, and were extracted with equal volumes of ethyl acetate three times. The ethyl acetate fractions containing the estrone-derived metabolites were pooled, vacuum-dried, and analyzed using UPLC–HRMS.

### Preparation of cell-extracts

Frozen strain DHT3 cell pellets (5 g) were suspended in 15 mL of HEPES-K^+^ buffer (pH 8.0; 100 mM). Strain DHT3 cells were broken by passing the cell suspensions through a French pressure cell (Thermo Fisher Scientific) twice at 137 megapascals. The cell lysates were fractionated using two steps of centrifugation steps. First, the cell-lysates were centrifuged at 20,000 × g for 30 min to get rid of most cell debris, unbroken cells and residual estradiol. Second, the supernatants containing the crude cell-extracts were centrifuged at 100,000 × g for 1.5 h to fractionate the soluble proteins from the membrane-bound proteins. All steps used for preparation of cell-extracts were performed at 4°C under anaerobic conditions.

### Cell-extract estradiol methylation activity assays

Assays were routinely performed in the dark (prevent abiotic cob(I)alamin oxidation) at 30°C for 1.5 h under an N_2_ atmosphere. The assay mixtures (1 mL) contained HEPES-K^+^ buffer (pH 8.0; 100 mM), soluble protein fraction (5 mg) extracted from strain DHT3 cells, estradiol (0.25 mM; [16,16,17-D3]estradiol and unlabeled estradiol in a 1:1 molar ratio), ATP (5 mM), MgSO_4_ (10 mM), and methylcobalamin (1 mM). In some assays, Ti(III) citrate-reduced cob(I)alamin (1 mM), SAM (0.25 mM), NADH (2 mM), and propyl-iodide (2.5 mM) were added to the assay mixtures. After 1.5 hours of incubation, the assays were extracted twice with the same volume of ethyl acetate. The ethyl acetate fractions containing estradiol-derived compounds were pooled, vacuum-dried, and stored at −20°C before further analytical analysis.

### Phylogenetic analysis

a. **cobalamin-dependent estradiol methyltransferase system EmtABCD.** The amino acid sequences of B9N43_10325 (*emtA*) from strain DHT3, *emtA* orthologs CBW56_16915 from *D. oestradiolicum* DSM 16959 and AMN47925.1 from *S. denitrificans* DSM 18526, and sequences of cobalamin-dependent methyltransferases searched with EmtA as query in UniProt (manually reviewed) were selected. The amino acid sequences of other cobalamin-dependent methyltransferases that have been functionally characterized (64) were also used for this phylogenetic analysis (**Table S3**). Fifty protein sequences were aligned using MUSCLE (65) in MEGA X (66) without truncation. The unrooted maximum likelihood tree was constructed using the LG model for amino acid substitution plus Gamma distribution rates (G) after the Model Test in MEGA X. The NNI for ML heuristic method and NJ/BioNJ for tree initiation were used for tree inference. Branch support was determined by bootstrapping 1,000 times. The maximum likelihood tree was then visualized in MEGA X.
b. **16S rRNA genes of selected steroid-degrading bacteria and their organization of genes involved in HIP catabolism.** The steroid-degrading bacteria with the corresponding sequenced genomes were chosen for this analysis. Protein sequences deduced from the previously characterized HIP degradation genes were used as queries to identify the orthologs in these genomes (**Table S6**). The deduced protein sequences from *Comamonas testosteroni* strain CNB-2 (a model organism for aerobic testosterone catabolism) and from *Mycobacterium tuberculosis* strain H37Rv (a model organism for aerobic cholesterol catabolism) were used as the queries for the Blastp analysis. Ortholog identification in each selected genomes was based on the criteria (e-value < 1e-5; pairwise identify ≥ 40%; query coverage ≥ 80%) of the Blastp results.

## ACKNOWLEDGMENTS

This research was funded by the Ministry of Science and Technology of Taiwan (MOST 107-2311-B-001-021-MY3). We gratefully appreciate Ms. Yu-Ching Wu at the Small Molecule Metabolomics Core Facility, Institute of Plant and Microbial Biology, Academia Sinica for the UPLC–HRMS analysis and Dr. Mei-Yeh Lu and her team at the High-Throughput Genomics Core Facility of the Biodiversity Research Center, Academia Sinica for genome sequencing. We also gratefully acknowledge Prof. Tony Z. Jia in Earth-Life Science Institute, Tokyo Institute of Technology for English editing. We also acknowledge Ms. Ching-Yen Tsai, Ms. I-Ting Lin, Mr. Yi-Shiang Huang and Mr. Yu-Sheng Wang for their contributions in culture enrichments, RNA-Seq and cell-extract activity assays.

## Notes

The authors declare no conflict of interest.

