## Supplemental Table S1-S5 and Figures S1-S8 for "Retroconversion of estrogens into androgens by bacteria *via* a cobalamin-mediated methylation"

### Supplementary Information

#### Table of Contents


##### Dataset

**Dataset S1.** Genome annotation of strain DHT3 and transcriptomic analysis (RNA-Seq) of bacterial cells grown anaerobically with testosterone or estradiol.

##### SI Tables

**Table S1.** Oligonucleotides used in this study.

**Table S2.** Selection of housekeeping genes of strain DHT3 used for constructing the linear regression line in the global gene expression profiles (RNA-Seq).

**Table S3.** Selection of the cobalamin-dependent methyltransferases used for the un-rooted maximum likelihood tree construction.

**Table S4.** UPLC–APCI–HRMS data of the intermediates involved in anaerobic estrone catabolism by strain DHT3.

**Table S5.** $^{1}$H- (600 MHz) and $^{13}$C-NMR (150 MHz) spectral data of the HPLC-purified metabolite (AND2**)** and the authentic standard 5α-androstan-3β,17β-diol

**Table S6.** Selection of the bacteria used for comparative analysis of the gene organization for HIP degradation.

##### SI Figures

**Fig. S1** Scanning electron micrographs of strain DHT3 cells.

**Fig. S2** Cobalamin as an essential vitamin during the anaerobic growth of strain DHT3 on estradiol.

**Fig. S3** Arrangement and expression analysis of the *emt* genes in strain DHT3.

**Fig. S4** The anaerobic growth of the wild type (A) and the *emtA*-disrupted mutant (B) of strain DHT3 with testosterone and estradiol.

**Fig. S5** APCI–HRMS spectrum of the HIP produced by estrone-fed strain DHT3.

**Fig. S6** UPLC–APCI–HRMS spectra of two TLC-purified androgen metabolites, 17β-hydroxyandrostan-3-one (A) and 3β,17β-dihydroxyandrostane (B).

**Fig. S7** SAM addition facilitates the estradiol methylation to form AND2 in the strain DHT3 cell-extracts.

**Fig. S8** The structures of the proteobacterial and actinobacterial gene clusters for the HIP catabolism.

### Appendix

**Appendix S1.** Nucleotide sequence of the 16S rRNA gene (B9N43_08010) of strain DHT3

**Appendix S2.** Nucleotide sequence of the *emtA* gene (B9N43_10325) of strain DHT3

**Appendix S3.** Nucleotide sequence of the *emtB* gene (B9N43_10320) of strain DHT3

**Appendix S4.** Nucleotide sequence of the *emtC* gene (B9N43_10315) of strain DHT3

**Appendix S5.** Nucleotide sequence of the *emtD* gene (B9N43_10310) of strain DHT3

### Dataset

**Dataset S1.** Genome annotation of strain DHT3 and transcriptomic analysis (RNA-Seq) of bacterial cells grown anaerobically with testosterone or estradiol (in a separated spread sheet).

### SI Tables

**Table S1.** Oligonucleotides used in this study.

| Primer | Sequence | Usage |
|---|---|---|
| 10305_emtD_1F | AGACGTTCCTTGCCTAGTGC | Operon analysis: intergenic region B9N43_10305/*emtD* |
| 10305_emtD_1R | GGAGGTTTTCCATCAGCCGA | |
| emtD_emtC_2F | GCCTATTTCAGCAATGGCCG | Operon analysis: intergenic region *emtD* /*emtC* |
| emtD_emtC_2R | ATCCTCTTCCCAGCTCTCGT | |
| emtC_emtB_3F | ACGAGAGCTGGGAAGAGGAT | Operon analysis: intergenic region *emtC*/*emtB* |
| emtC_emtB_3R | TTTGACTGCACCGATCACGA | |
| emtB_emtA_4F | AAAAAGCGGTCGAGCACAAC | Operon analysis: intergenic region *emtB* /*emtA* |
| emtB_emtA_4R | AGCAGGTCCACCGTAGTTTG | |
| emtA_emt10330_5F | AACCTGGGACTGCTATGCAC | Operon analysis: intergenic region *emtA*/ B9N43_10330 |
| emtA_emt10330_5R | GAACCCGGGGACTTGCATAA | |
| 10330_10335_6F | CGCCATTCAATACGGGCTTG | Operon analysis: intergenic region B9N43_10330/_10335 |
| 10330_10335_6R | TGTCCACCAGTTCGTCGAAG | |
| emtA_F | ATGATTCCAAGCATTGATTTCCAGC | *emtA*-specific primers (expected product size = 1.4 kb) |
| emtA_R | TTAGCGATAGGGCATGCC | |
| emtA_773\|774s IBS | AAAAAAGCTTATAATTATCC TTACTGGACGAGTTCGTGCG CCCAGATAGGGTG | TargeTron Gene Knockout system: *emtA*-specific primer |
| emtA_773\|774s EBS1d | CAGATTGTACAAATGTGGTG ATAACAGATAAGTCGAGTTC GATAACTTACCTTTCTTTGT | TargeTron Gene Knockout system: *emtA*-specific primer |
| emtA_773\|774s EBS2 | TGAACGCAAGTTTCTAATTTC GGTTTCCAGTCGATAGAGGAA AGTGTCT | TargeTron Gene Knockout system: *emtA*-specific primer |
| EBS Universal | CGAAATTAGAAACTTGCGTTCA GTAAAC | TargeTron Gene Knockout system: universal primer |

**Table S2.** Selection of housekeeping genes of strain DHT3 used for constructing the linear regression line in the global gene expression profiles (RNA-Seq) in **Figure 2B**. Strain DHT3 cells were anaerobically grown with estradiol or testosterone.

| **Locus tag** | **Gene name** | **Definition** | **Testosterone [log2(X+1)]** | **Estradiol [log2(X+1)]** |
|---|---|---|---|---|
| B9N43_01505 | *aceE* | pyruvate dehydrogenase (acetyl-transferring), homodimeric type | 8.05 | 9.25 |
| B9N43_03845 | *gltA* | citrate (Si)-synthase | 6.42 | 6.94 |
| B9N43_03880 | - | malate dehydrogenase | 8.92 | 9.56 |
| B9N43_04270 | - | pyruvate dehydrogenase (acetyl-transferring) E1 component subunit alpha | 2.16 | 4.21 |
| B9N43_07665 | - | transcription termination factor Rho | 8.27 | 10.62 |
| B9N43_08640 | - | DNA gyrase subunit A | 8.96 | 9.81 |
| B9N43_09345 | *ftsZ* | cell division protein FtsZ | 8.82 | 10.57 |
| B9N43_09365 | - | preprotein translocase subunit SecA | 9.91 | 10.89 |
| B9N43_11395 | - | RNA polymerase sigma factor RpoD | 11.09 | 10.58 |
| B9N43_12040 | *recA* | recombinase RecA | 10.95 | 10.57 |
| B9N43_12290 | *gyrB* | DNA topoisomerase (ATP-hydrolyzing) subunit B | 6.69 | 8.43 |
| B9N43_12535 | *glnA* | type I glutamate--ammonia ligase | 7.27 | 9.35 |
| B9N43_15425 | *gap* | type I glyceraldehyde-3-phosphate dehydrogenase | 9.22 | 10.05 |
| B9N43_15595 | - | isocitrate dehydrogenase (NADP(+)) | 9.03 | 9.69 |
| B9N43_15770 | *tuf* | elongation factor Tu | 10.43 | 10.26 |
| B9N43_15810 | *rpoB* | DNA-directed RNA polymerase subunit beta | 5.95 | 7.80 |
| B9N43_15815 | *rpoC* | DNA-directed RNA polymerase subunit beta' | 7.88 | 9.60 |
| B9N43_15830 | *fusA* | elongation factor G | 6.11 | 7.30 |
| B9N43_16390 | *gap* | type I glyceraldehyde-3-phosphate dehydrogenase | 4.33 | 3.91 |

**Table S3.** Selection of the cobalamin-dependent methyltransferases used for the un-rooted maximum likelihood tree construction. The functions of listed proteins were manually reviewed (Swiss-Port) or experimentally characterized except AMN47925.1.

| Enzyme | Subunit | Microorganism | Taxonomy (Class) | Accession No. |
|---|---|---|---|---|
| **Estradiol methyltransferases from denitrifying proteotacteria** | | | | |
| estradiol methyltransferase | | *Denitratisoma* sp. strain DHT3 | Betaproteobacteria | This study |
| estradiol methyltransferase | | *Denitratisoma oestradiolicum* DSM 16959 | Betaproteobacteria | This study |
| estradiol methyltransferase | | *Steroidobacter denitrificans* DSM 18526 | Gammaproteobacteria | GeneBank AMN47925.1 |
| **Methyltetrahydrofolate:homocysteine methyltransferases (methionine synthases) from bacteria** | | | | |
| Methionine synthase (Group 2) | MetH | *Escherichia coli* K12 | Gammaproteobacteria | UniPort P13009 |
| Methionine synthase (Group 1) | | *Mycobacterium tuberculosis* ATCC 25618 | Actinobacteria | UniPort O33259 |
| Methionine synthase (Group 2) | | *Salmonella typhimurium* ATCC 700720 | Gammaproteobacteria | UniPort P37586 |
| Methionine synthase (Group 2) | | *Vibrio cholerae* serotype O1 ATCC 39315 | Gammaproteobacteria | UniPort Q9KUW9 |
| Methionine synthase (Group 1) | | *Mycobacterium leprae* TN | Actinobacteria | UniPort Q49775 |
| Methionine synthase (Group 2) | | *Pseudomonas aeruginosa* ATCC 15692 | Gammaproteobacteria | UniPort Q9I2Q2 |
| Methionine synthase (Group 1) | | *Synechocystis* sp. PCC 6803 | Cyanophyceae | UniPort Q55786 |
| Methionine synthase (Group 2) | | *Aliivibrio fischeri* ATCC 700601 | Gammaproteobacteria | UniPort Q5E814 |
| Methionine synthase (Group 2) | | *Vibrio parahaemolyticus* serotype O3:K6 RIMD 2210633 | Gammaproteobacteria | UniPort Q87L95 |
| Methionine synthase (Group 2) | | *Vibrio vulnificus* YJ016 | Gammaproteobacteria | UniPort Q7MHB1 |
| Methionine synthase (Group 2) | | *Vibrio vulnificus* CMCP6 | Gammaproteobacteria | UniPort Q7MHB1 |
| Methionine synthase (Group 2) | | *Aliivibrio fischeri* | Gammaproteobacteria | UniPort Q9AJQ8 |
| **Coenzyme M (CoM) methyltransferases from methanogenic archaea** | | | | |
| Monomethylamine:CoM methyltransferase | MtmB | *Methanosarcina barkeri* | Methanomicrobia | UniPort O30642 |
| | | *Methanosarcina mazei* DSM 3647 | Methanomicrobia | UniPort P58969 |
| | | *Methanosarcina barkeri* DSM 804 | Methanomicrobia | UniPort P0C0W4 |
| | | *Methanosarcina barkeri* | Methanomicrobia | UniPort Q9P9L4 |
| | | *Methanosarcina acetivorans* ATCC 35395 | Methanomicrobia | UniPort P58865 |
| | | *Methanosarcina acetivorans* ATCC 35395 | Methanomicrobia | UniPort P58866 |
| | | *Methanosarcina barkeri* DSM 804 | Methanomicrobia | UniPort Q46E72 |
| | | *Methanosarcina barkeri* DSM 804 | Methanomicrobia | UniPort P0C0W3 |
| Dimethylamine:CoM methyltransferase | MtbB | *Methanosarcina barkeri* DSM 804 | Methanomicrobia | UniPort P0C0W5 |
| | | *Methanosarcina barkeri* DSM 804 | Methanomicrobia | UniPort P0C0W6 |
| | | *Methanosarcina barkeri* | Methanomicrobia | UniPort O93661 |
| | | *Methanosarcina mazei* DSM 3647 | Methanomicrobia | UniPort P58971 |
| | | *Methanosarcina mazei* DSM 3647 | Methanomicrobia | UniPort P58970 |
| | | *Methanosarcina mazei* DSM 3647 | Methanomicrobia | UniPort P58972 |
| | | *Methanosarcina barkeri* | Methanomicrobia | UniPort Q9P9N0 |
| | | *Methanosarcina acetivorans* ATCC 35395 | Methanomicrobia | UniPort Q8TS72 |
| | | *Methanosarcina acetivorans* ATCC 35395 | Methanomicrobia | UniPort Q8TN68 |
| | | *Methanosarcina acetivorans* ATCC 35395 | Methanomicrobia | UniPort Q8TTA5 |
| | | *Methanosarcina barkeri* | Methanomicrobia | UniPort Q9P9M9 |
| Trimethylamine:CoM methyltransferase | MttB | *Methanosarcina barkeri* | Methanomicrobia | UniPort O93658 |
| | | *Methanosarcina acetivorans* ATCC 35395 | Methanomicrobia | UniPort Q8TTA9 |
| | | *Methanosarcina* mazei DSM 3647 | Methanomicrobia | UniPort P58973 |
| | | *Methanosarcina mazei* DSM 3647 | Methanomicrobia | UniPort P58974 |
| | | *Methanosarcina thermophila* | Methanomicrobia | UniPort Q9P995 |
| | | *Methanosarcina barkeri* DSM 804 | Methanomicrobia | UniPort P0C0W7 |
| | | *Methanosarcina acetivorans* ATCC 35395 | Methanomicrobia | UniPort Q8TS73 |
| | | *Methanococcoides burtonii* DSM 6242 | Methanomicrobia | UniPort Q12TR2 |
| Methanol:CoM methyltransferase | MtaB | *Methanosarcina barkeri* DSM 804 | Methanomicrobia | UniPort Q46EH3 |
| Dimethylsulfide:CoM methyltransferase | MtsB | *Methanosarcina barkeri* | Methanomicrobia | UniPort Q48925 |
| | | *Methanosarcina mazei* DSM 3647 | Methanomicrobia | UniPort Q8PUA7 |
| **Tetrahydrofolate ( THF) methyltransferases from acetogenic or methylotrophic bacteria** | | | | |
| Veratrol:THF *O*-demethylase | OdmB | *Acetobacterium dehalogenans* | Clostridia | GeneBank AAC83696.2 |
| Vanillate:THF *O*-methyltransferase | MtvB | *Moorella thermoacetica* ATCC 39073 | Clostridia | GeneBank YP429263.1 |
| glycine betaine:THF methyltransferase | MtgB | *Desulfitobacterium hafniense* Y51 | Clostridia | UniPort Q24SP7 |
| Acetyl-CoA synthase/CO dehydrogenase | AcsE | *Moorella thermoacetica* | Clostridia | UniPort Q46389 |
| Chloromethane:THF methyltransferase | CmuA | *Methylobacterium extorquens* CM4 | Alphaproteobacteria | GeneBank CAB39403.1 |

**Table S4.** UPLC–HRMS data of the intermediates involved in anaerobic estrone catabolism by strain DHT3. The steroid substrate was composed of unlabeled estrone and [3,4C-$^{13}$C]estrone (mixed in a 1:1 molar ratio). The bacterial culture was extracted using ethyl acetate, and the metabolites were analyzed through UPLC–APCI–HRMS. MS data shown are from the unlabeled metabolites.

| Compound ID | UPLC behavior (RT[a], min) | Molecular formula (predicted molecular mass)[b] | Dominant ion peaks | Identification of product ions |
|---|---|---|---|---|
| testosterone | 6.01 | $C_{19}H_{28}O_2$<br>288.21 | 289.21 | $[M+H]^+$ |
| androst-4-en-3,17-dione | 5.74 | $C_{19}H_{26}O_2$<br>286.19 | 287.19 | $[M+H]^+$ |
| DT | 5.57 | $C_{19}H_{26}O_2$<br>286.19 | 287.20 | $[M+H]^+$ |
| androsta-1,4-diene-3,17-dione | 5.30 | $C_{19}H_{24}O_2$<br>284.18 | 285.18 | $[M+H]^+$ |
| 1-testosterone | 6.32 | $C_{19}H_{28}O_2$<br>288.21 | 289.21 | $[M+H]^+$ |
| 17-hydroxy-androstan-1,3-dione | 5.15 | $C_{19}H_{28}O_3$<br>304.21 | 287.20<br>305.21 | $[M-H_2O+H]^+$<br>$[M+H]^+$ |
| 2,3-SAOA | 5.08 | $C_{19}H_{30}O_4$<br>322.22 | 305.21<br>323.22 | $[M-H_2O+H]^+$<br>$[M+H]^+$ |
| HIP | 2.39 | $C_{13}H_{18}O_4$<br>238.12 | 221.12<br>239.13 | $[M-H_2O+H]^+$<br>$[M+H]^+$ |

[a]RT, retention time. [b]The predicated molecular mass was calculated using the atom mass of $^{12}C$ (12.0000), $^{16}O$ (15.9949), and $^{1}H$ (1.0078).

**Table S5.** $^1$H- (600 MHz) and $^{13}$C-NMR (150 MHz) spectral data of the HPLC-purified androgen metabolite (AND2**)** and the authentic standard 5α-androstan-3β,17β-diol (=3β,17β-dihydroxyandrostane) purchased from Steraloids, Inc.

| Positions | AND2 | | | 5α-androstan-3β,17β-diol | |
|---|---|---|---|---|---|
| | $^1$H $^{a,\,b}$ | $^{13}$C | | $^1$H $^{a,\,b}$ | $^{13}$C |
| 3 | 3.51 (1H, m) | 72.0 | | 3.50 (1H, m) | 72.0 |
| 17 | 3.55 (1H, t, *J* = 8.7) | 82.7 | | 3.55 (1H, t, *J* = 8.7) | 82.7 |
| 18 | 0.85 (3H, s) | 11.8 | | 0.85 (3H, s) | 11.8 |
| 19 | 0.72 (3H, s) | 12.9 | | 0.72 (3H, s) | 12.9 |

[a] Measured in methanol-$d_4$; [b] δ in ppm, mult. (*J* in Hz).

**Table S6.** Selection of the bacteria used for comparative analysis of the gene organization for HIP degradation in **Fig. 6** (in a separated spreadsheet).

### SI Figures

**Fig. S1** Scanning electron micrographs of the strain DHT3 cells.

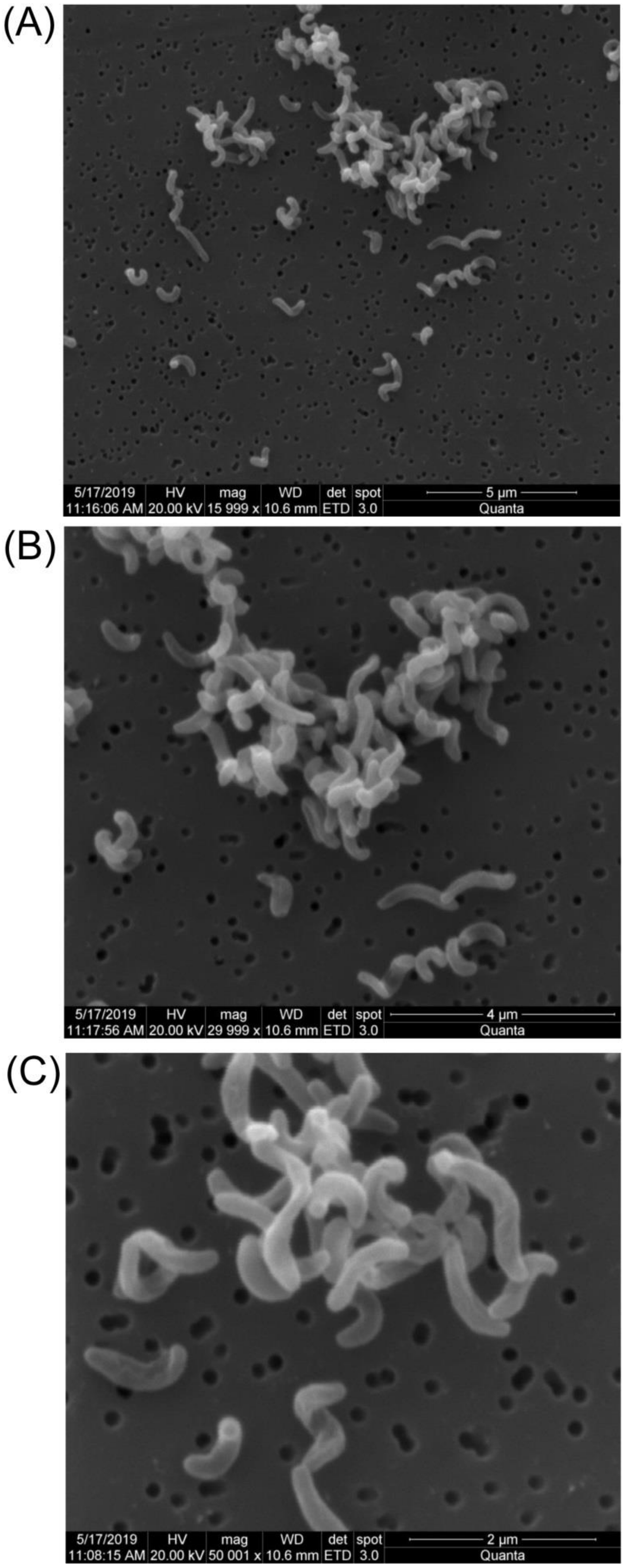

**Fig. S2** Cobalamin as an essential vitamin during the anaerobic growth of strain DHT3 on estradiol. The fed-batch cultures (100 mL) of strain DHT3 were treated with or without vitamins. In all of the treatments, estradiol (3 mM) served as the sole carbon and energy source, whereas nitrate (initially 10 mM) served as terminal electron acceptor. Bacterial growth was measured as the total protein concentration in the cultures, and the data shown are averages (deviations <5%) of three experimental measurements.

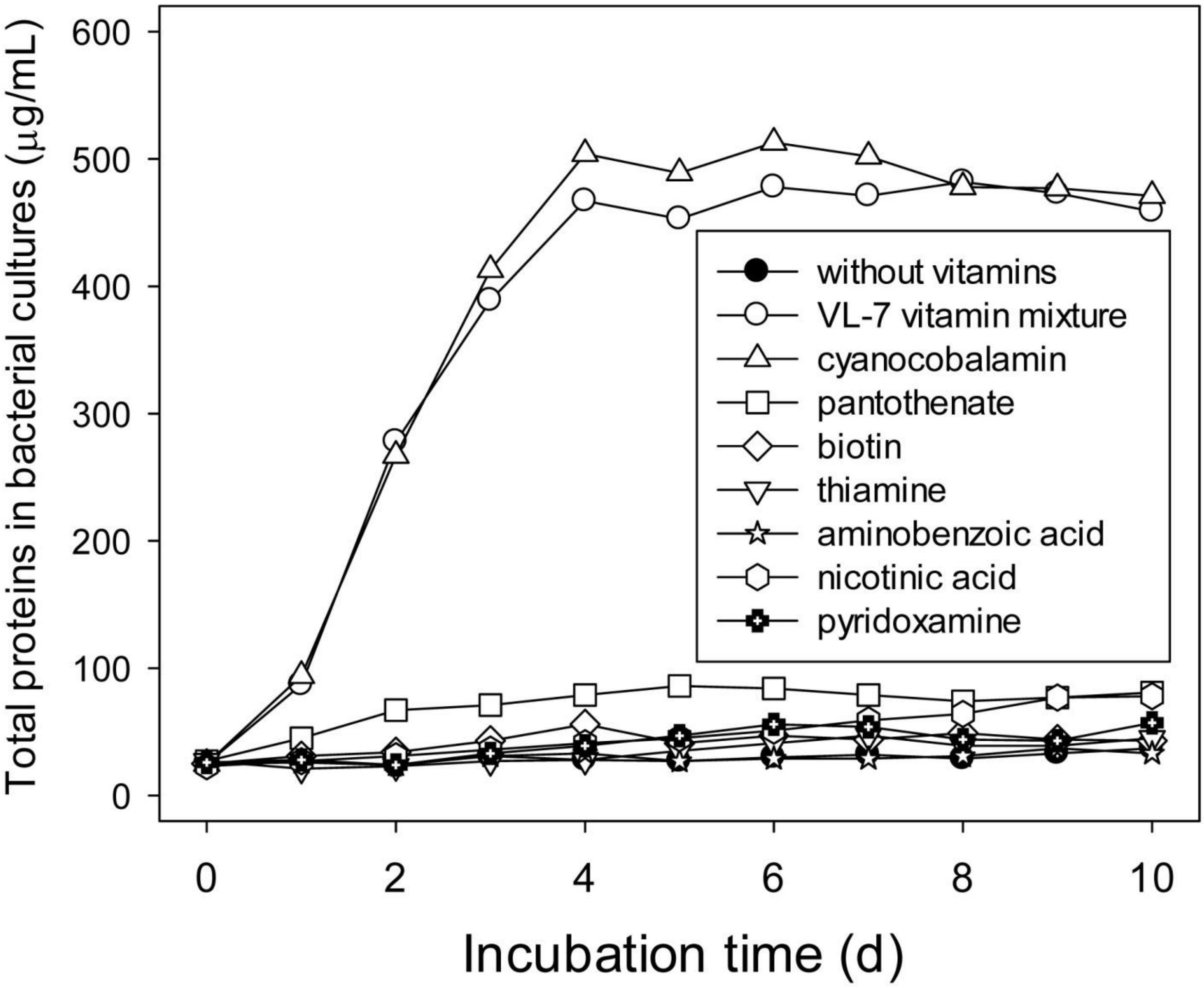

**Fig. S3** Arrangement and expression analysis of the *emt* (**e**stradiol **met**hylation) genes in strain DHT3. The *emt* operon of strain DHT3 comprises five genes (B9N43_10310~10330). For operon analysis, total RNA from strain DHT3 was extracted and the contaminating DNA was removed. cDNA was synthesized by reverse transcriptase using random hexamers as primers. To analyze the transcriptional unit, cDNA was used as template for PCR to bridge the intergenic regions of the *emt* genes. Each primer set was applied to 3 different PCR templates, including genomic DNA (lane 1), cDNA synthesized using random hexamers (lane 2), and total RNA (lane 3). Genomic DNA and total RNA were used as positive and negative controls, respectively. Oligonucleotides used in this study are listed in **Table S1** and their location is indicated by black arrows.

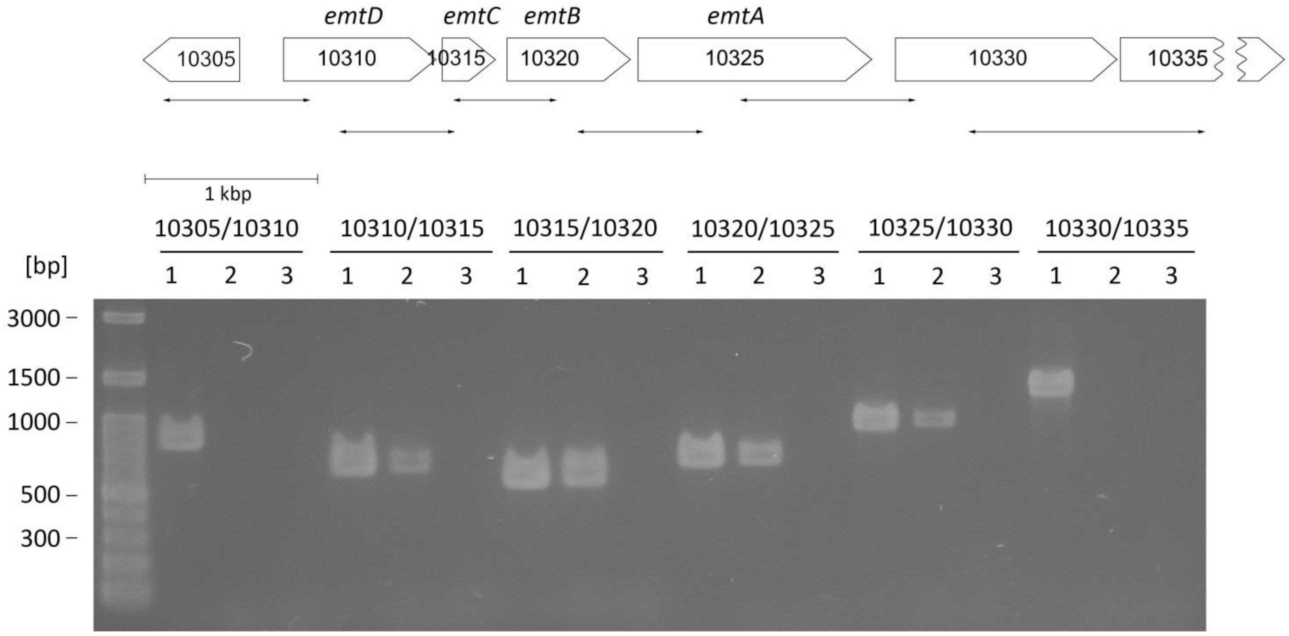

**Fig. S4** The anaerobic growth of the wild type and the *emtA*-disrupted mutant of strain DHT3 with testosterone (A) and estradiol (B).

(A)

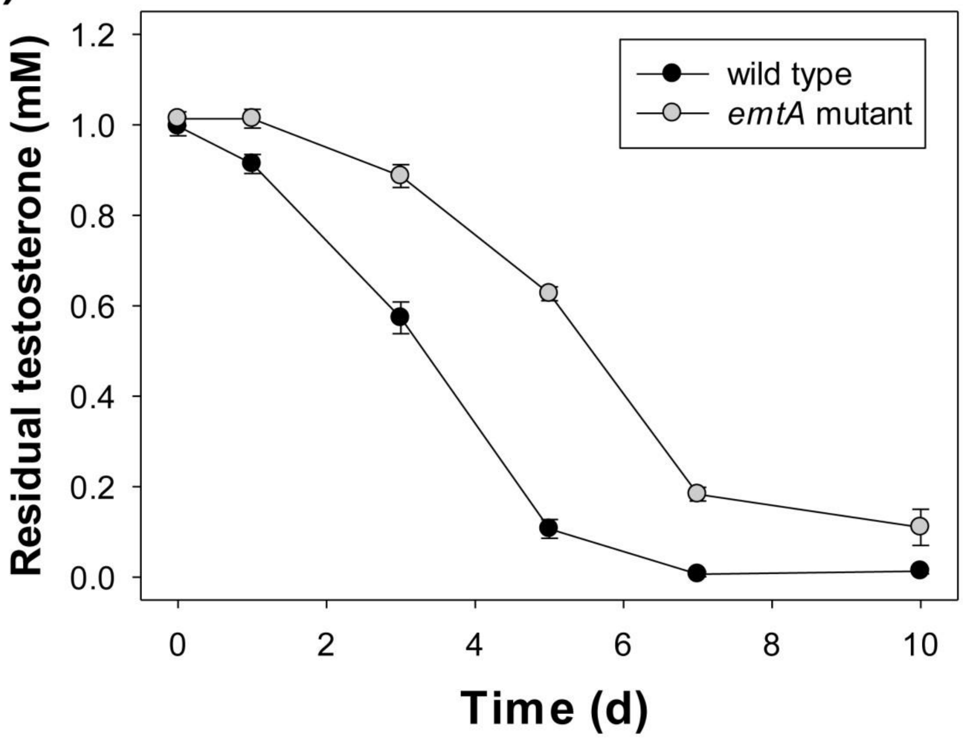


(B)

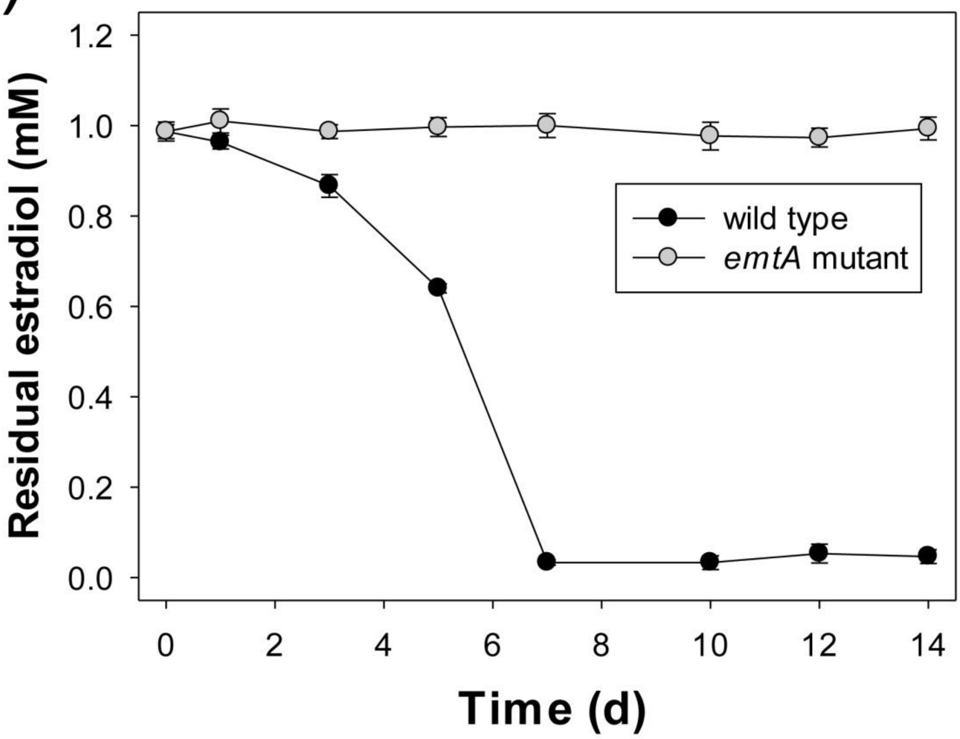

**Fig. S5** APCI–HRMS spectrum of the HIP produced by the estrone-fed strain DHT3. The steroid substrate was composed of unlabeled estrone and [3,4C-$^{13}$C]estrone (mixed in a 1:1 molar ratio). The bacterial culture was extracted using ethyl acetate, and the metabolites were analyzed through UPLC–APCI–HRMS.

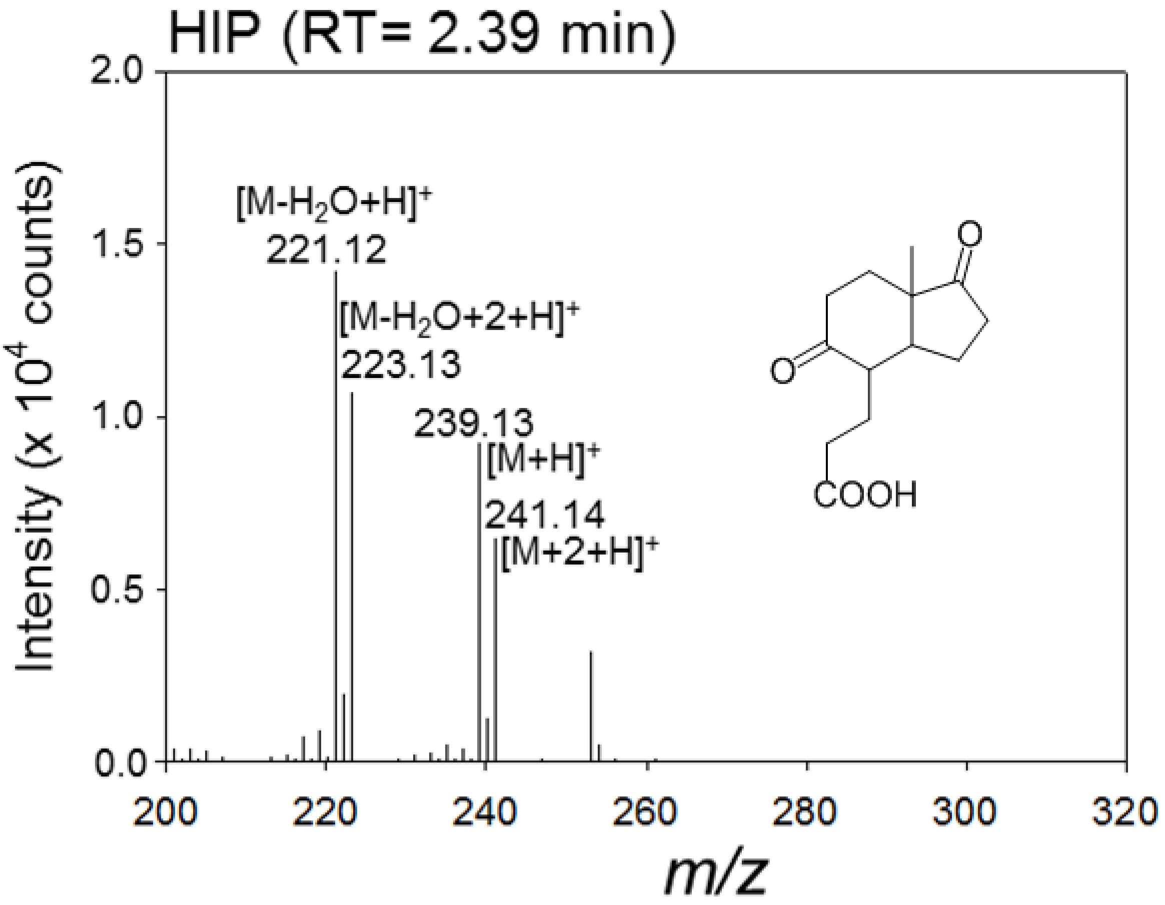

**Fig. S6** UPLC–APCI–HRMS spectra of two TLC-purified androgen metabolites, 17β-hydroxyandrostan-3-one (A) and 3β,17β-dihydroxyandrostane (B). The androgens were produced by the strain DHT3 cell extract incubated with estradiol ([16,16,17-D3]17β-estradiol and unlabeled estradiol in a 1:1 molar ratio). See **Fig. 6Bi1** for the chemical composition of the reaction mixture. The predicted elemental composition of individual intermediates was calculated using Xcalibur™ Software Mass Spectrometry Software (Thermo Fisher Scientific).

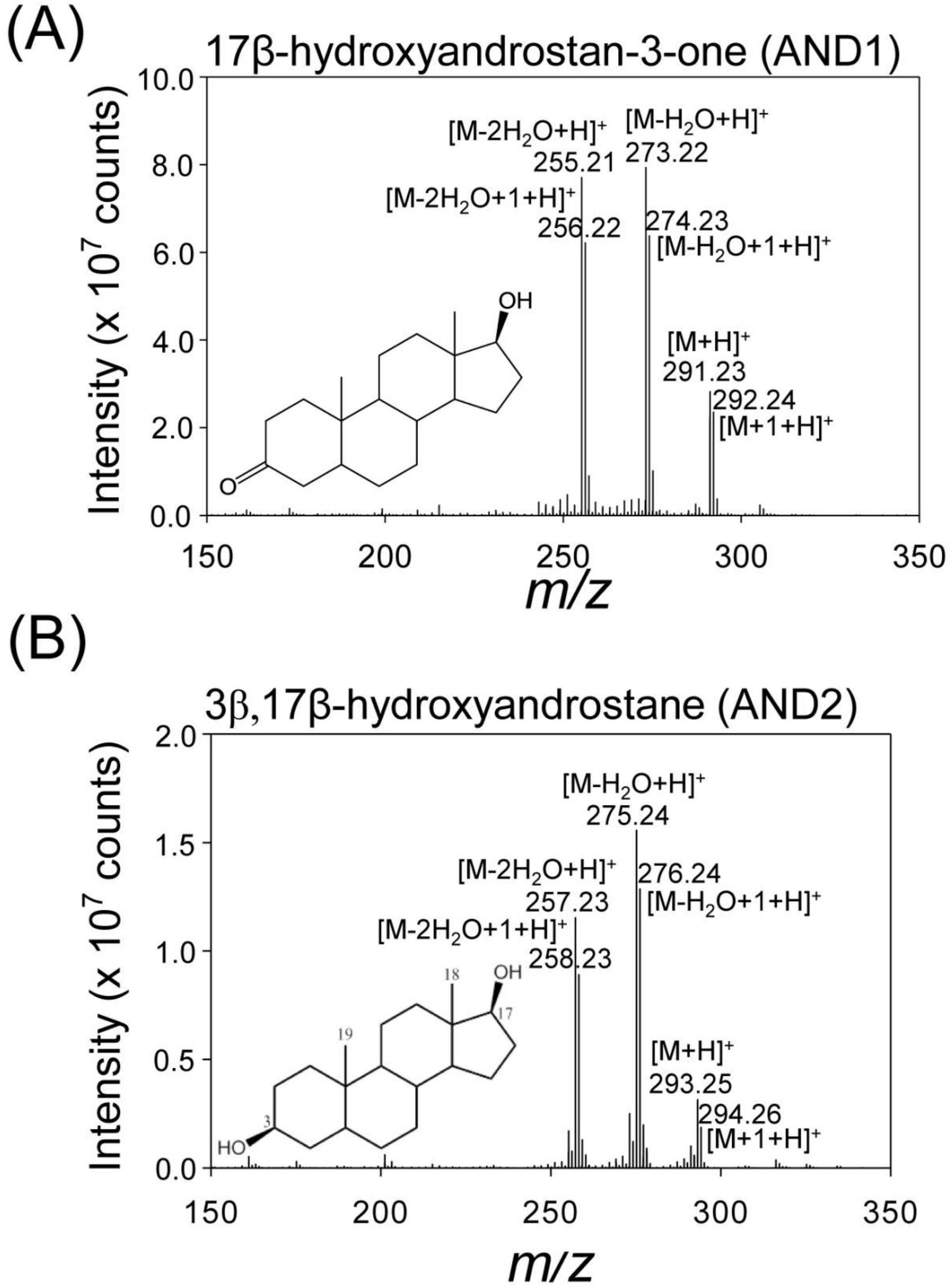

**Fig. S7** SAM addition facilitates the estradiol methylation to form AND2 in the strain DHT3 cell-extracts. The androgen production exhibited a dose-dependent manner with SAM addition. The assay mixtures (1 mL) contained 100 mM HEPES-$K^+$ buffer (pH 8.0), strain DHT3 cell-extracts (5 mg), estradiol (0.25 mM), ATP (5 mM), $MgSO_4$ (10 mM), NADH (2 mM), SAM (0–0.25 mM), and with or without propyl iodide (2.5 mM). Abbreviations: E2, 17β-estradiol.

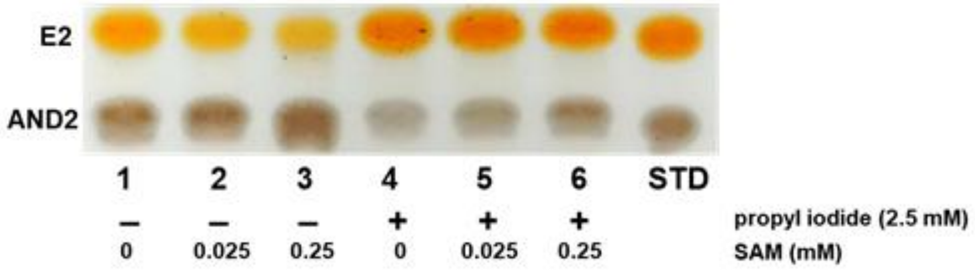

**Fig. S8** Phylogenetic tree of the selected steroid-degrading bacteria and the distinguishable organization of HIP degradation genes in proteobacteria and actinobacteria (see **Table S6** for the detailed information of the selected bacterial strains). The phylogenetic trees were constructed using the neighbor-joining method with Jukes–Cantor parameter and a bootstrap value of 1000. Bacterial strains in bold: the steroid degradation capabilities have been experimentally verified. Bacterial strains in red: able to anaerobically degrade steroids.

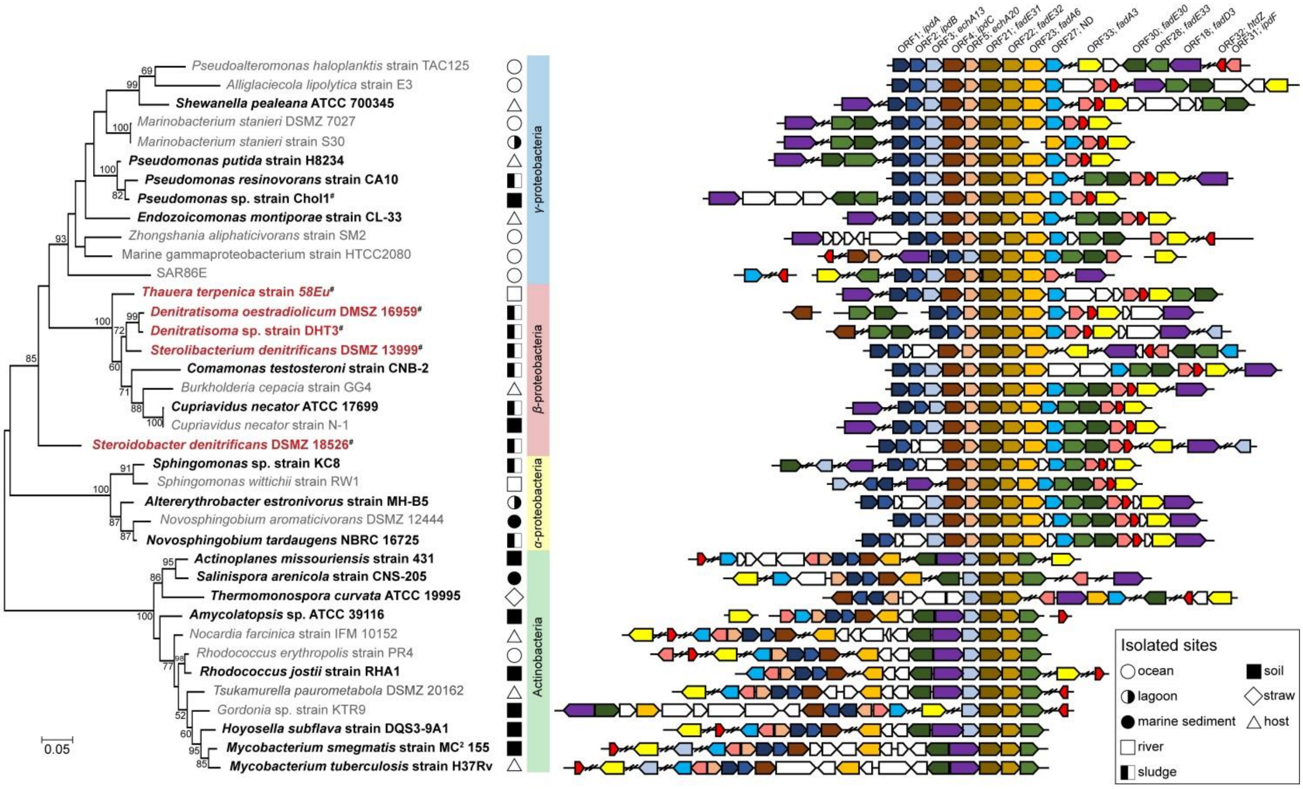

### Appendix

**Appendix S1.** Nucleotide sequence of the 16S rRNA gene (B9N43_08010) of strain DHT3.

AGAGATTGAACTGAAGAGTTTGATCCTGGCTCAGATTGAACGCTGGCGGAATGCTTTACA
CATGCAAGTCGAACGGCAGCACGGGTGCTTGCACCTGGTGGCGAGTGGCGAACGGGTG
AGTAATACATCGGAACGTGCCCAGTAGAGGGGGATAACCGTCCGAAAGGATGGCTAATAC
CGCATACGCCCTGAGGGGGAAAGCGGGGGACCGCAAGGCCTCGCGTTATTGGAGCGGCC
GATGTCGGATTAGCTAGTTGGTGGGGTAAAGGCCTACCAAGGCGACGATCCGTAGCTGGT
CTGAGAGGATGATCAGCCACACTGGGACTGAGACACGGCCCAGACTCCTACGGGAGGCA
GCAGTGGGGAATTTTGGACAATGGGGGCAACCCTGATCCAGCCATTCCGCGTGAGTGAA
GAAGGCCTTCGGGTTGTAAAGCTCTTTCGGCAGGAACGAAAAGGTGGTCACTAATACTG
GCTACTGATGACGGTACCTGAAGAAGAAGCACCGGCTAACTACGTGCCAGCAGCCGCGG
TAATACGTAGGGTGCGAGCGTTAATCGGAATTACTGGGCGTAAAGCGTGCGCAGGCGGTT
TTGTAAGACAGCTGTGAAATCCCCGGGCTTAACCTGGGAACTGCGGTTGTGACTGCAAG
ACTGGAGTGTGGCAGAGGGGGGTGGAATTCCACGTGTAGCAGTGAAATGCGTAGAGATG
TGGAGGAACACCGATGGCGAAGGCAGCCCCCTGGGTTAACACTGACGCTCATGCACGAA
AGCGTGGGGAGCAAACAGGATTAGATACCCTGGTAGTCCACGCCCTAAACGATGCCAACT
AGGTGTTGGGGAAGGAGACTTCCTTAGTACCGTAGCTAACGCGTGAAGTTGGCCGCCTG
GGGAGTACGGTCGCAAGATTAAAACTCAAAGGAATTGACGGGGACCCGCACAAGCGGT
GGATGATGTGGATTAATTCGATGCAACGCGAAAAACCTTACCTACCCTTGACATGCCAGG
AACTTTCCAGAGATGGATTGGTGCCCGAAAGGGAGCCTGGACACAGGTGCTGCATGGCT
GTCGTCAGCTCGTGTCGTGAGATGTTGGGTTAAGTCCCGCAACGAGCGCAACCCTTGTCA
TTAGTTGCCATCATTTAGTTGGGCACTCTAATGAGACTGCCGGTGACAAACCGGAGGAAG
GTGGGGATGACGTCAAGTCCTCATGGCCCTTATGGGTAGGGCTTCACACGTCATACAATG
GTCGGTACAGAGGGTTGCCAAGCCGCGAGGTGGAGCCAATCCCAGAAAGCCGATCGTAG
TCCGGATTGGAGTCTGCAACTCGACTCCATGAAGTCGGAATCGCTAGTAATCGCGGATCA
GCATGTCGCGGTGAATACGTTCCCGGGTCTTGTACACACCGCCCGTCACACCATGGGAGT
GGGTTTCACCAGAAGTAGGTAGTCTAACCGCAAGGAGGGCGCTTACCACGGTGGGGTTC
ATGACTGGGGTGAAGTCGTAACAAGGTAGCCGTATCGGAAGGTGCGGCTGGATCACCTC
CTTTCTA

**Appendix S2.** Nucleotide sequence of the *emtA* gene (B9N43_10325) of strain DHT3.

ATGATTCCAAGCATTGATTTCCAGCGGCGCTCAACCACCGGACCTGTTGGCAAGACCGAC
GACTTTGACCTGGATCTGGCCTTCAAGGTCAGGGAACTGGTGGAGACCTACAACATCAA
GTACGACCCCAACCAACTTGTGGTGGATGACCGCACGGCAGATGCAATTTTTGATGCGGG
CGTCGAATTGCTGGCCGAGGTCGGCCTGTTTCATCAGCAGACCTCGCGCATCATGCTGTA
CTCCAAGGAGGAACTCTATCAACTGGCGGCCGAGTCCAAGGCGAACCCAGCCTGCATTC
CCTTTGGAAAAGGCGAGGACCGGATGTATCTGCGGCATCGCAAGAGTACAGACACCTTT
GCCCCGACAAACTACGGTGGACCTGCTGGAGTCGCTGAACCGGAGTGGTTCATTCCCTAT
GTTCAGTCCTTTGCCCAGGAGCGTCATGTAAAGGGCCTGGGGATTTGTCCGGGCGTACCG
CGCATTGGCGATCTTGACCCGAAGGCCGGTACCCTGACCGAAGTGGAAATCGCCCTTTGG
GAGCAGGAAGCCCTGCGTGAGGCGCTGAAGCGCACTGGCCGTCTGAACATGAACCTGG
GACTGCTATGCACCGCCAGCACCCCCTCGGGCACCATGTCGGTCATGGCCAGTGGGTATC
GGGACCATCTGAATACCCAGATCGGCATTCACATCATGCCCGAACAGAAAATGAGCTGGA
ATCCGCTGCTGCTTTCCCAGTACTGCGAGAACGCCGGGATCGAGCCCTGGATGAGTTCGA
TGTCCTGCATCGGCGGTTTGTGTCGGGATGCCGCCGAGGTCGCAGTAACAATGGTGGCTA
ACGCGTTGGGGCAACTCAGCTATGCCAAGGGCGGCTCCATGAGCTACTTCCCCAGCCATC
TGGATGGGACATGGGCAACGCGACCTTCTCATTGGGCATTCAGTGGCGCAGCCCGTGCTT
CTGAACGTCACCTTGGACTGGCTGTGGGAACCTCTATCTCGGGCATCACCAATGCCTGGC
GCACTCCCTTGACCCTGTGGCAGTCGGCGGCGGTCGTGTTGACCTCCGTGGCCAGCGGA
TTGTCCTATGCCTGGATTTCCGGACATACCGGCCTGGAGGCGCGACTGATCGGAGAAATG
ATGGATGTTTGCGCTGGCATGCCGGCAAAGGAAGCCAATGAACTGGCCCAGCGGGTTAT
GGTCAAGGTTGATGAGCTTCTGCCTCAGGTAACGAAGCAGTTGCCCTTCGTCGAAGCTTA
TGACATCGAAACCGTTCAGCCCCGTCCGGCATACGAGTCTTCCATGCTGAAGGTGCGGGA
TGAACTGCAGCGTATGGGCATGCCCTATCGCTAA

**Appendix S3.** Nucleotide sequence of the *emtB* gene (B9N43_10320) of strain DHT3

ATGAGCAGCATTGAGGCAATCCGTGAGGCCGTCTGGCGCCTGAAGAAAAAGGATGCCGT
TGCCCTGGTGGAAGAAGGCCTTGCAGAAGGGCTTGATCCTACTGCGATGTTGAAGGAAG
GGGTCATTGCCGGTTTGCAGGAGGTCGGGCGCAAGTTTGGTGCCGGCGAGTATTTTCTGG
CAGAACTGGTGATGGCTGGCAAGGTCGGTGAGCCCTGTATCGACTTGATCACGCCTCACT
TACCGCCGAATTCGGAAGGGAAGATGGGAACGGTCGTGATCGGTGCAGTCAAAGGCGAT
CTGCACACCATTGGCTACGGGCTGGTGACAACCCAACTGGAGTTGGCGGGATTCGAAGT
CATCAAGCTGGGCATCGACCTGGACAGCAAGTACTTCATCGAAAAAGCGGTCGAGCACA
ACGCCGACATCATCGGCCTGTCCGCCTTCCTGGTGACGACCATTCCCTATTGCCCAGAAG
TCTTGGGTTATTTGAAGGACATGGGGCTGCGGGATCGCTTCAAGGTCATCATCGGCGGCA
CGGAATGTACCGCCGACAAGGCGGACGCCATGGATGCCGATGGCTGGGCACTCAATGCA
ATCTCAGCGGTGCCACTGTGCAAGCGCCTGATGGGCAAGGACGTGGGTGAGGAGGCCAA
GCTCGCGCAAACCTACGACCACGGCTGGTGGTACAACTCCCGCCGCCATGGCACCTGA

**Appendix S4.** Nucleotide sequence of the *emtC* gene (B9N43_10315) of strain DHT3

ATGAAGCTCGAAGAATTAAAAACGCAGTTCCCGGATCTGACATTTGAGGAGGTCGGTCCT
GACGAGAGCTGGGAAGAGGATGGCGCCGGATATTGCTTCGTTATAGAGGCGATGATCAAG
GACAAGCGTGTTCGCGCTACCGTCATGGTGGATGACTTGCCGCGGGTGGCCTTCACGGGT
GACTCTGTGGGCATCCGCAAGATGCTGGTGAGGTGGGCAGTGGGGCGGGATCACTTGGC
ATATCTGGCCTATGAGCTGGGCCGGGCCGAGATGGCGATCCGTCACGGCGTCCCCTTCCT
TCAGGAGTGA

**Appendix S5.** Nucleotide sequence of the *emtD* gene (B9N43_10310) of strain DHT3

ATGTTCATGCAGTTCTGGCAATCACTGTTTCTGACGGAAGGGGACCAGATACTTGATGTG
ACCAAGATATGCGATGGTCTTGGTTTCCACGGGATGCTGTTTCCCGATCATCTGATCCACC
CCGAGAAGCAGGACTCCACCTACCTGTACTCGGCTGATGGAAAACCTCCATCATTTACGG
AGGACACAGTATGGCCTGAGTGTTGGTCGCTATTTGCGACGTTGGCGGCTATGACCAAGA
ATCTGCATTTCTGCACATGCGTTTTCATTCTTCCCTTACGCAATCCGATTGAACTGGCCAA
GGCAACGTCAAGCGTTGCCTATTTCAGCAATGGCCGTATCCATCTGGGCGCCGGGGCCGG
CTGGATGAAGGAAGAGTTCGAGATGCTCGGCGTGGATTGGGCCACGCGCGGAAAGCGAT
ATGACGAGTGTATCGAGGTGATGCGCAAGCTCTATACGGGGCAGTACGTCGAACACCACG
GCGAATTCTTCGATTTCCCACGCATCATGATGACGCCGGTTCCAGAGAAACCCGTGCCCA
TCCTGATCGGCGGCATCAGCGGCCCCGCCTTGCGTCGAGCTGCCCGTATTGGCGATGGTT
GGATCGGGCCGGGACAGAGTGTCGATGCGGCCCTGCAAACCCTGAGTACCCTGAATAAG
TTGCGGACCGAGTACGGAACGCAGAACAAGGAATTCAACAACATCGTTCCTATCTACGG
GGATGTCAGCATCGACGACATCAAGCGACTGGAGGACGCTGGGGCTACAGGGATGGTCA
GCTTGCCCTTTGCTTTCACAATCAAACCAGGAACGACCTTGGAGGAAAAGCGTGCCTATC
TGGAGCGCTATTCCCAGGAAGTCATTGCGAAATTCAGGTGA
